# Dynein directs prophase centrosome migration to control the stem cell division axis in the developing *C. elegans* epidermis

**DOI:** 10.1101/2023.09.25.559295

**Authors:** Cátia Carvalho, Daniel J. Barbosa, Ricardo Celestino, Esther Zanin, Ana Xavier Carvalho, Reto Gassmann

## Abstract

The microtubule motor dynein is critical for the assembly and positioning of mitotic spindles. In *C. elegans*, these dynein functions have been extensively studied in the early embryo but remain poorly explored in other developmental contexts. Here we use a hypomorphic dynein mutant to investigate the motor’s contribution to asymmetric stem cell-like divisions in the larval epidermis. Live imaging of seam cell divisions that precede formation of the seam syncytium shows that mutant cells properly assemble but frequently mis-orient their spindle. Mis-oriented divisions misplace daughter cells from the seam cell row, generate anucleate compartments due to aberrant cytokinesis, and disrupt asymmetric cell fate inheritance. Consequently, the seam becomes disorganized and populated with extra cells that have lost seam identity, leading to fatal epidermal rupture. We show that dynein orients the spindle through the cortical GOA-1^Gαi^–LIN-5^NuMA^ pathway, which directs the migration of prophase centrosomes along the anterior–posterior axis. Spindle mis-orientation in the dynein mutant can be rescued by stretching cells, implying that dynein–dependent cortical cues and elongated cell shape jointly ensure correct asymmetric division of epithelial stem cells.

## INTRODUCTION

Animal mitosis requires the assembly of a microtubule-based chromosome segregation machine called the spindle, in which microtubules grow out from centrosome-containing spindle poles (Valdez *et al*., 2023). Astral spindle microtubules, which in most cells make contact with the cell cortex, are used to position the spindle (di Pietro *et al*., 2016; McNally, 2013), and this in turn determines the division axis and the relative size of daughter cells. In oriented divisions, which play a major role in the development and maintenance of tissues (Lechler and Mapelli, 2021), the spindle axis aligns with a pre-defined axis in the tissue. Regulated positioning of the spindle is particularly important for asymmetric cell division that generates daughter cells with distinct fates, such as the choice between self-renewal versus differentiation in stem cells (Bergstralh and St Johnston, 2014; Dewey *et al*., 2015; Williams and Fuchs, 2013).

Among the mechanisms that control spindle positioning are cortical cues. These typically involve the microtubule minus-end-directed motor complex cytoplasmic dynein 1 (dynein), which contains two copies each of a motor heavy chain (HC), an intermediate chain (IC), a light intermediate chain, and three light chains. When localized at the cell cortex together with its obligatory co-factor dynactin, dynein can exert pulling forces on astral microtubules, and studies in *C. elegans*, *D. melanogaster*, and human cells have identified a set of core components required for this activity (Kotak, 2019; Kotak and Gönczy, 2013; Kiyomitsu, 2019): the G protein subunit Gαi, which has two *C. elegans* orthologues called GOA-1 and GPA-16, is anchored in the plasma membrane and in its GDP–bound form interacts with C-terminal GoLoco motifs in *D. melanogaster* Pins, *C. elegans* GPR1/2 (G-protein regulators 1 and 2) and vertebrate LGN (leucine glycine asparagine). N-terminal tetricopeptide repeats in these proteins then recruit the coiled-coil adaptor for dynein–dynactin, called *D. melanogaster* Mud (Mushroom body defective), *C. elegans* LIN-5 (abnormal cell lineage 5) and vertebrate NuMA (nuclear mitotic apparatus).

In *C. elegans*, studies addressing dynein–dependent cortical force generation during asymmetric cell division have focused on the P-cell lineage in the early embryo (Colombo *et al*., 2003; Couwenbergs *et al*., 2007; Gotta and Ahringer, 2001; Grill *et al*, 2001; Grill *et al*., 2003; *Gusnowski et al.*, 2011; Miller and Rand, 2000; Nguyen-Ngoc *et al*., 2007; Portegijs *et al*., 2016; Rodriguez-Garcia *et al*., 2018; Srinivasan *et al*., 2003; Tsou *et al*., 2003). This has revealed instances in which the direction of dynein–dependent centrosome or spindle movement correlates with asymmetric distribution of the motor’s cortical recruitment machinery. In the embryonic founder cell P0, sequential enrichment of GPR-1/2 at the anterior cortex during prophase and the posterior cortex at metaphase/anaphase is thought be at least partly responsible for the asymmetry of pulling forces that center the nuclear–centrosome complex and displace the spindle toward the posterior, respectively (Colombo *et al*., 2003; Gotta *et al*., 2003; Park and Rose, 2008). In the P2 cell of the four–cell embryo, GPR-1/2 become enriched at the EMS/P2 boundary, which likely helps bring about the rotation of the nuclear–centrosome complex in late prophase such that one of the nascent spindle poles faces the GPR-1/2–enriched region (Heppert *et al*., 2018; Werts *et al*., 2011; Zhang *et al*., 2008).

In addition to positioning the spindle, dynein plays essential roles in spindle assembly, including nuclear attachment of centrosomes and their separation to opposite sides of the nucleus in prophase (Raaijmakers and Medema, 2014). In the *C. elegans* early embryo, dynein inhibition prevents prophase centrosome separation and, consequently, formation of a bipolar spindle, resulting in chromosome segregation failure (Gönczy *et al*., 1999; Schmidt *et al*., 2005). Dynein’s essential role during embryonic cell division makes it challenging to examine how the motor contributes to asymmetric cell divisions during post-embryonic development. Furthermore, dynein’s involvement in spindle assembly means that dynein *null* mutants are of limited use when trying to assess the motor’s specific contribution to spindle orientation.

Seam cells in the *C. elegans* epidermis are a model for asymmetric division within a polarized epithelium (Chisholm and Hsiao, 2012). In larvae, seam cells form a linear row on each lateral side. Apical junctions connect seam cells with each other and with the surrounding hyp7 syncytium, which is already generated in the embryo. Larval development of the seam involves a stereotypical series of proliferative (L2 stage) and self-renewing asymmetric divisions (L1, L2, and L3 stage) that expand the number of seam cells and generate daughters with distinct fates, respectively. Following an asymmetric division, the anterior daughter cell undergoes terminal differentiation either by fusing with hyp7 or adopting a neuronal identity. The posterior daughter retains seam identity and goes on to form an apical junction with its neighbor, thereby closing the gap in the seam left by the differentiated anterior daughter. Seam cells can therefore be regarded as stem cells with limited potential for self-renewal. At the L4 stage, the 16 lateral seam cells fuse with each other to generate the mature seam syncytium. Among other functions within the epidermis, the seam secretes components of the cuticle, the external structure that acts as a permeability barrier and is critical for the integrity of the animal (Chisholm and Xu, 2012).

The Wnt/β–catenin asymmetry pathway, a variant of canonical Wnt signaling, plays important roles during seam cell division. Its components, which include the APC homolog APR-1, are asymmetrically distributed along the anterior–posterior axis (Mizumoto and Sawa, 2007), and this is critical for correct cell fate of daughter cells (Gleason and Eisenmann, 2010; Yamamoto *et al*., 2011). Additionally, the Wnt/β–catenin asymmetry pathway controls the division axis by providing cues for orienting the mitotic spindle along the anterior–posterior axis. This function is redundant with geometrical cues provided by cell shape (Wildwater *et al*., 2011), which remains elongated in mitosis due to tension propagated within the seam cell row through apical junctions. The role of dynein during seam development, including the motor’s potential contribution to oriented division, has not been investigated.

Here, we describe a *C. elegans* dynein mutant that results in a disorganized seam. Using live imaging, we find that the mutant perturbs spindle orientation, but not spindle assembly, during the last round of asymmetric seam cell divisions. This hypomorphic phenotype allowed us to assess the importance of oriented division for seam development, including cell fate decisions. We present the first characterization of the cortical dynein pathway in seam cells, which shows how dynein ensures spindle orientation along the anterior–posterior axis and reveals interplay between dynein and cell shape-based cues.

## RESULTS

### An N-terminal deletion mutant of *C. elegans* dynein IC supports development but results in premature death of adults by epidermal rupture

In an effort to determine the functional significance of the dynein IC N-terminus, which binds the dynein regulators dynactin p150 and Nde1, we used CRISPR/Cas9-mediated genome editing to delete residues 4 to 36 in the *C. elegans* dynein IC homolog DYCI-1 (**Fig. 1A; Fig. S1A**). Animals homozygous for the mutation, hereafter referred to as *dyci-1(11N)*, develop into fertile adults, which lay embryos that fail to hatch. This contrasts with animals homozygous for the null allele *dyci-1(tm4732)*, which become developmentally arrested between larval stages L2 and L3 (**Fig. 1A, B**). These findings indicate that dynein maintains residual activity when the interactions mediated by the DYCI-1 N-terminus are missing, and that this residual activity is sufficient to support development to adulthood and fertility. The hypomorphic nature of *dyci-1(11N)* therefore offers an opportunity to examine the roles of dynein at developmental stages that are inaccessible with dynein null mutants.

**Figure 1:**
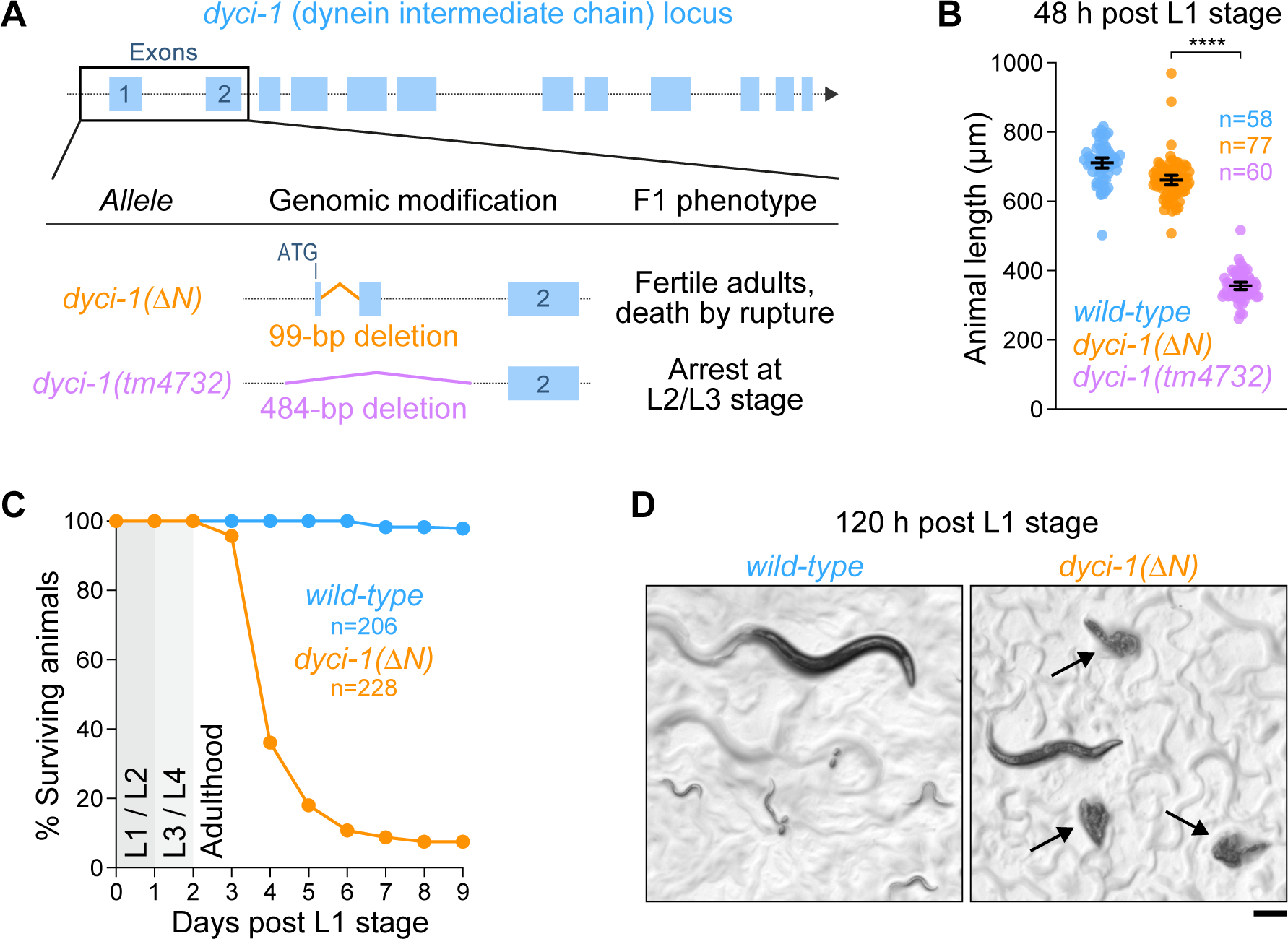
An N-terminal deletion mutant of *C. elegans* dynein intermediate chain supports development but results in premature death of adults by epidermal rupture. **(A)** Schematic of the genomic *dyci-1* locus and the two deletion mutants examined in this study. The in-frame deletion *dyci-1(11N)* results in dynein intermediate chain lacking residues 4 - 36, whereas the more extensive region deleted in *dyci-1(tm4732)* includes the start codon. **(B)** Animal length measured 48 h after release from L1 arrest, showing the contrasting effects of *dyci-1(tm4732)* versus *dyci-1(11N)* on development. Each plotted data point corresponds to a measurement of an individual animal, and *n* denotes the number of animals examined. Bars show the mean ± 95% CI. Statistical significance was determined by the Mann-Whitney test. \*\*\**\*P* < 0.0001. **(C)** Lifespan analysis during the first 9 days after release from L1 arrest. *n* denotes the number of animals examined. **(D)** Images of animals on an NGM plate seeded with bacteria 120 h after release from L1 arrest. Arrows point to adult *dyci-1(11N)* animals that died due to spontaneous epidermal rupture, which is responsible for the rapid decline in adult viability shown in *(C)*. Scale bar, 0.2 mm.

Surprisingly, we found that developmentally synchronized *dyci-1(11N)* animals started to die 3 days after being released from L1 arrest, and 90 % of animals were dead 3 days later (**Fig. 1C**). *dyci-1(11N)* animals therefore die shortly after reaching adulthood. Closer inspection revealed that *dyci-1(11N)* animals overwhelmingly (92 %, n=212) died by rupturing (**Fig. 1D**). Dynein is required for proper formation of the vulva (Celestino *et al*., 2019), the weakest point in the cuticle, but differential interference contrast imaging showed that *dyci-1(11N)* animals did not rupture through the vulva (**Fig. S1B**). We conclude that *dyci-1(11N)* weakens cuticle integrity beyond the vulva, causing penetrant premature death by spontaneous epidermal rupture.

### The *dyci-1(ΔN)* mutation compromises epidermal seam integrity

The penetrant rupture phenotype of *dyci-1(11N)* animals suggested that the mutation compromises cuticle integrity. Consistent with this idea, the cuticle of *dyci-1(11N)* animals was permeable to the normally excluded dye Hoechst 33342 when assayed 72 h after release from L1 arrest (**Fig. 2A, B**). Cuticle formation requires secretion from epidermal cells (Chisholm and Xu, 2012). To test whether the cuticle integrity defect in *dyci-1(11N)* animals reflects compromised dynein function in the epidermis, we expressed wild-type *dyci-1* from a single-copy integrated transgene under the epidermal promoter P*dpy-7*. Epidermal expression of transgenic wild-type *dyci-1* in *dyci-1(11N)* animals rescued cuticle permeability and prevented premature death by rupture (**Fig. 2C**). By contrast, *dyci-1(11N)* animals expressing transgenic *dyci-1(11N)* in the epidermis still ruptured (72 % of dead animals, n=158). These results show that dynein function in the epidermis is required for cuticle integrity.

**Figure 2:**
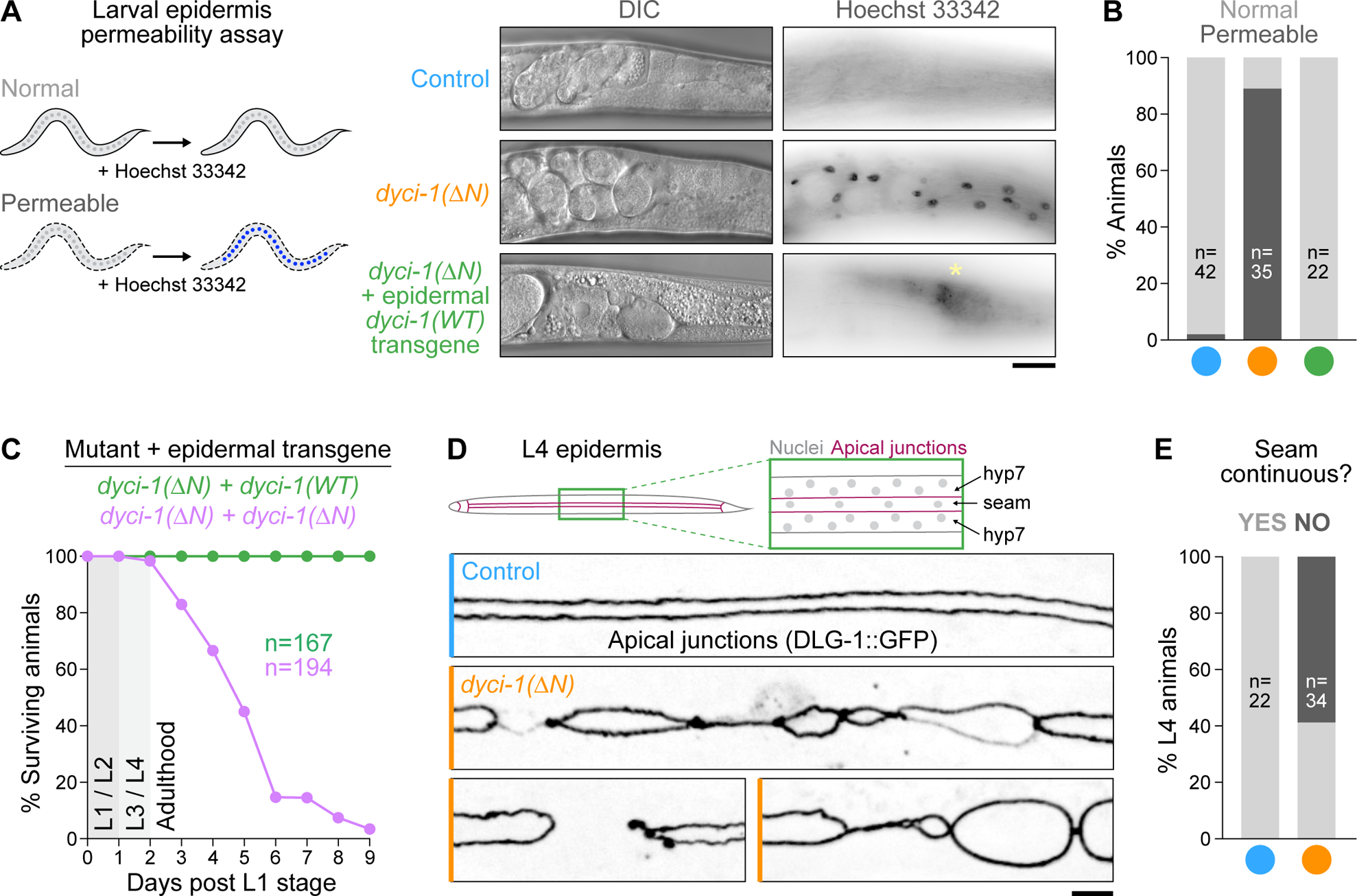
The *dyci-1(ΔN)* mutation compromises epidermal seam integrity. **(A)** *(left)* Schematic illustrating that the membrane–permeable dye Hoechst 33342 stains epidermal nuclei only if the barrier function of the external surface (the cuticle) is compromised. *(right)* Differential interference contrast (DIC) and corresponding fluorescence images showing Hoechst 33342 staining of epidermal nuclei in adult *dyci-1(11N)* animals, which is rescued by expression of transgene-encoded DYCI-1 in the epidermis. The asterisk marks autofluorescence signal from the intestine in the image of the transgenic animal. Scale bar, 30 µm. **(B)** Quantification of epidermal permeability using the assay shown in *(A)*. *n* denotes the number of animals examined. **(C)** Lifespan analysis during the first 9 days after release from L1 arrest, showing that the rescue of epidermal permeability in *(A, B)* correlates with survival. *n* denotes the number of animals examined. **(D)** (top) Schematic showing the location of apical junctions in the late-L4 epidermis, which connect the syncytial hyp7 cell with the lateral seam syncytium. *(bottom)* Fluorescence images of late-L4 animals expressing DLG-1::GFP, a marker for apical junctions, showing seam discontinuities in the *dyci-1(11N)* mutant. **(E)** Quantification of seam discontinuities based on imaging as shown in *(D)*. *n* denotes the number of animals examined.

To examine epidermal tissue morphology in *dyci-1(11N)* animals, we imaged the apical junction component DLG-1::GFP. In control L4 animals, DLG-1::GFP localized to two continuous parallel lines per lateral side, which mark the border between the seam syncytium and epidermal cells, primarily the large syncytial hyp7 cell (**Fig. 2D, E**). In *dyci-1(11N)* animals, the seam was frequently disrupted by gaps and compartments of variable sizes. A similarly disorganized seam syncytium, as well as animal death by rupture, had previously been described for co-inhibition of the ninein-like protein NOCA-1 and the patronin homolog PTRN-1 (Wang *et al*., 2015). Since co-inhibition of NOCA-1 and PTRN-1 perturbs the circumferential microtubule array in the hyp7 cell (Wang *et al*., 2015), we asked whether aberrant seam morphology in *dyci-1(11N)* animals could be caused by a disorganized microtubule cytoskeleton. Visualization of epidermal microtubules with GFP::β-tubulin (TBB-2) showed that the *dyci-1(11N)* mutation, in contrast to co-inhibition of NOCA-1 and PTRN-1, did not perturb the regular arrangement of microtubule bundles in hyp7 (**Fig. S1C, D**). We conclude that *dyci-1(11N)* causes severe morphological defects in the seam without affecting epidermal microtubule organization.

### The *dyci-1(ΔN)* mutation causes spindle mis-orientation during asymmetric seam cell division

To understand why *dyci-1(11N)* results in a disorganized seam, we monitored the last round of seam cell divisions at the L3 larval stage by time-lapse imaging in animals co-expressing GFP::TBB-2 and mCherry (mCh)-tagged histone H2B (HIS-11) (**Fig. 3A**). Control cells proceeded from nuclear envelope breakdown (NEBD) to anaphase onset in 6.2 ± 0.5 min (mean ± 95 % CI) (**Fig. 3B**), and divided along the A–P axis defined by the seam cell row. *dyci-1(11N)* cells assembled a bipolar spindle of normal size, which elongated in anaphase at the same rate as control spindles (**Fig. 3C, D**). Chromosome congression took slightly longer in *dyci-1(11N)* cells, which extended the NEBD–anaphase onset interval by 5 min on average (**Fig. 3B**), but there was no discernible chromosome mis-segregation in anaphase. In contrast to control cells, metaphase spindles in *dyci-1(11N)* were occasionally severely mis-oriented (**Fig. 3A**). Measurement of the angle between the spindle axis and the A–P (long) axis of cells in 3D image stacks revealed that spindle angles at anaphase onset were significantly larger in *dyci-1(11N)* cells relative to controls (**Fig. 3E**). We conclude that dynein is required for proper spindle orientation in seam cells.

**Figure 3:**
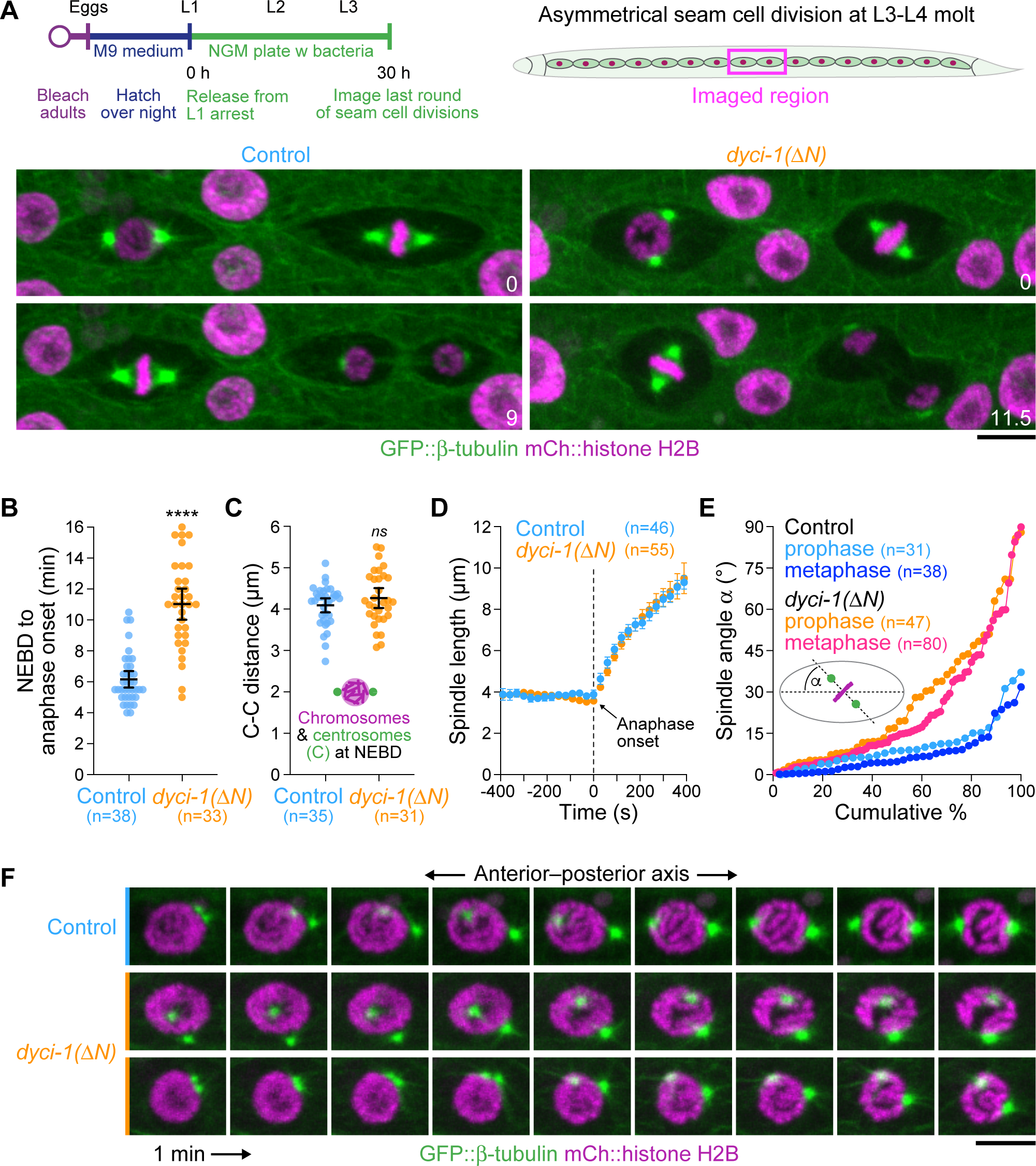
The *dyci-1(ΔN)* mutation results in spindle mis-orientation during asymmetric seam cell division. **(A)** *(top left)* Schematic of the experimental procedure followed for live imaging experiments. Animals are first synchronized at the L1 stage before release from the L1 arrest for 30 h. This time point is ideal for imaging the last round of seam cell divisions at the late L3 stage. *(top right)* Schematic showing the epidermal region imaged in subsequent experiments, which corresponds to dividing cells in the middle region of the seam cell row. *(bottom)* Selected images from time-lapse movies of dividing seam cells co-expressing GFP::TBB-2 (β-tubulin) and mCh::HIS-11 (histone H2B). Time is indicated in minutes. Scale bar, 5 µm. **(B) – (E)** Quantification of the interval between NEBD and anaphase onset *(B)*, centrosome– centrosome distance at the end of prophase *(C)*, spindle length over time *(D)*, as well as the angle of the centrosome–centrosome axis relative to the anterior–posterior axis at the end of prophase and just prior to anaphase onset *(E)*. Data points plotted in *(B)*, *(C)*, and *(E)* correspond to measurements in individual cells, and bars in *(B)* – *(D)* denote the mean ± 95% CI. In all graphs, *n* denotes the total number of cells imaged in at least 10 animals. Statistical significance was determined by the Mann-Whitney test. \*\*\**\*P* < 0.0001; *ns* = not significant, *P* > 0.05. **(F)** Successive frames (1 min apart) from time-lapse movies of cells as in *(A)*, showing that prophase centrosomes in the *dyci-1(11N)* mutant fail to separate along the anterior–posterior axis. Two examples are shown for the *dyci-1(11N)* mutant. Scale bar, 5 µm.

### The *dyci-1(ΔN)* mutation perturbs positioning of prophase centrosomes along the A–P axis

To better understand how dynein contributes to spindle orientation, we examined prophase centrosome separation. In control cells, centrosomes marked by GFP::TBB-2 separated along the A–P axis, one centrosome remaining relatively stationary while the other moved to the opposite side of the mCh::HIS-11–marked nucleus (**Fig. 3F**). *dyci-1(11N)* did not inhibit centrosome separation *per se* but instead affected the path of centrosome migration such that the centrosome–centrosome axis was poorly aligned with the A–P axis at NEBD (**Fig. 3C, F**). Quantitative analysis showed that centrosome–centrosome axis angles at NEBD were comparable to spindle axis angles at anaphase onset (**Fig. 3E**). These results suggest that dynein acts in early prophase to direct centrosome migration along the A–P axis and that the orientation of the spindle is set up prior to NEBD by alignment of the centrosome–centrosome axis with the A–P axis.

### Dynein acts at the seam cell cortex downstream of LIN-5^NuMA^

The observation that dynein contributes to spindle orientation in seam cells predicts that the motor is anchored at the cell cortex to generate pulling forces on astral microtubules. To test this, we examined the localization of dynein in larval L3-stage seam cells using endogenous GFP-tagged dynein HC (DHC-1), co-expressed with mCh::HIS-11 and the mCh-tagged pleckstrin homology (PH) domain of mammalian PLC1δ1 as a plasma membrane marker. During interphase and prophase, GFP::DHC-1 was highly concentrated near the apical junctions that connect neighboring seam cells and also exhibited a comparatively modest enrichment across the remaining regions of the cell cortex (**Fig. 4A, B**). GFP::DHC-1 also localized to mitotic centrosomes, spindle microtubules, and kinetochores, and became prominently enriched in the cleavage furrow at later stages of cytokinesis (**Fig. 4A**). In the *dyci-1(11N)* mutant, GFP::DHC-1 was no longer noticeably enriched at subcellular sites except for residual kinetochore signal (**Fig. S2A**). Diffuse cytoplasmic GFP::DHC-1 levels were increased in *dyci-1(11N)* cells relative to controls (**Fig. S2B**), suggesting the motor is de-localized in the mutant rather than degraded, consistent with immunoblotting results (**Fig. S1A**).

**Figure 4:**
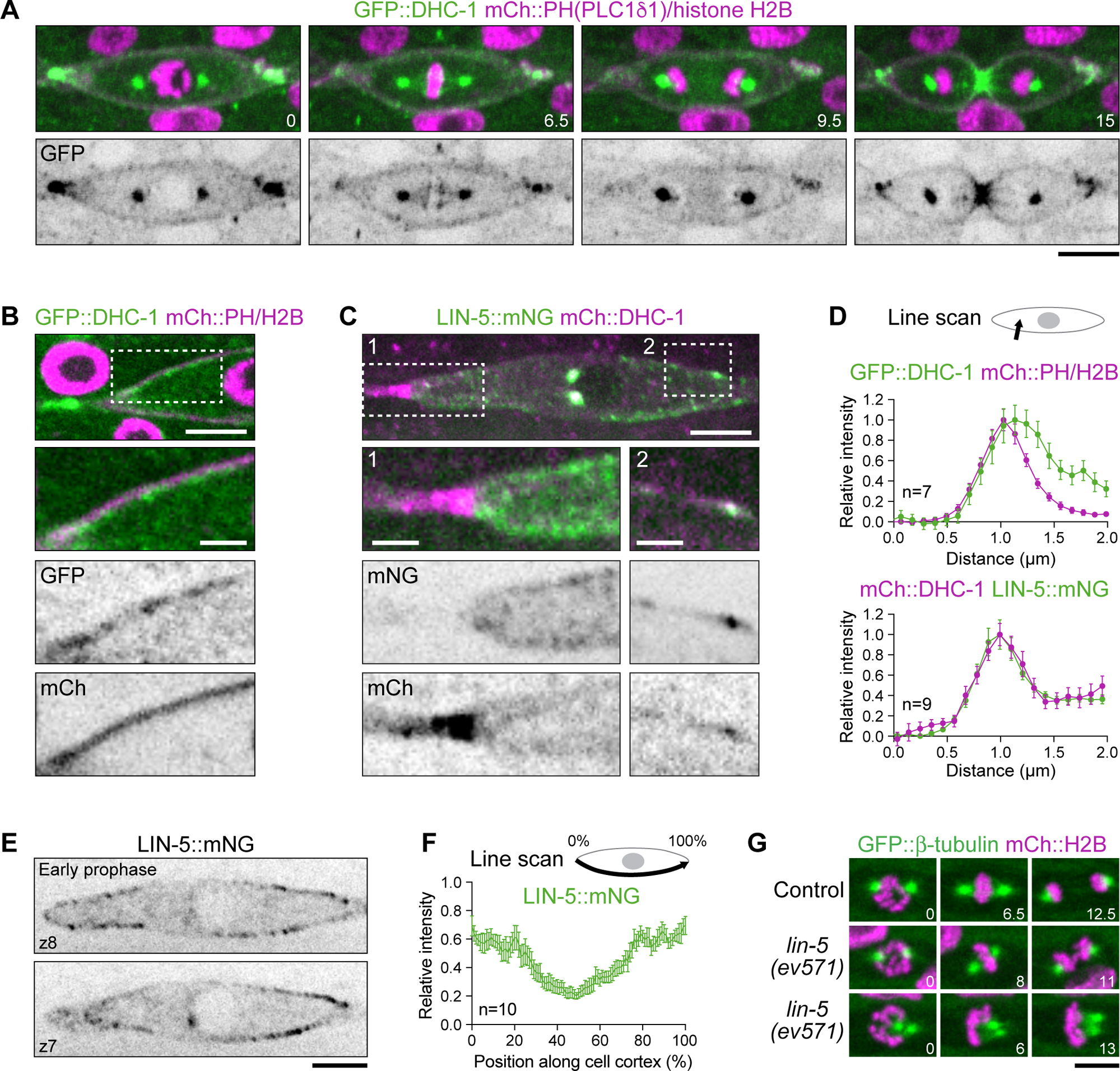
Dynein acts at the seam cell cortex downstream of LIN-5^NuMA^. **(A)** Selected images from a time-lapse movie in a dividing seam cell co-expressing GFP::DHC-1 (dynein heavy chain), the plasma membrane marker mCh::PH(PLC181), and mCh::HIS-11 (histone H2B). Time is indicated in minutes. Scale bar, 5 µm. **(B)** Image of a cortical region in a prophase seam cell as in *(A)*, showing enrichment of GFP::DHC-1 at the plasma membrane. Scale bars, 5 µm *(top)* and 2 µm *(magnified region)*. **(C)** Image of a prophase seam cell co-expressing LIN-5::mNG and mCh::DHC-1. Enlarged regions show partial overlap between the two markers at the cell cortex near, but not at, the junctions that connect neighboring seam cells. Scale bars, 5 µm *(top)* and 2 µm *(magnified region)*. **(D)** Line scan profiles across the seam cell cortex, as illustrated in the schematic (*top*), measured in images such as in *(B)* and *(C)*. Data is plotted as mean ± SEM. *n* denotes the total number of cells examined in at least 5 animals. **(E)** Single confocal z-section (z) images (0.5 µm apart) of the prophase seam cell in *(C)*, showing uneven cortical enrichment of LIN-5::mNG. Scale bar, 5 µm. **(F)** Line scan profile of LIN-5::mNG along the early prophase cell cortex, as indicated in the schematic *(top)*, in cells such as in *(E)*, showing that LIN-5::mNG distribution is bipolar. Data is plotted as mean ± SEM. *n* denotes the number of cells examined (one cell per animal). **(G)** Selected images from time-lapse movies of dividing seam cells co-expressing GFP::TBB-2 (β-tubulin) and mCh::HIS-11 (histone H2B), showing mitotic defects in the temperature-sensitive *lin-5(ev571)* mutant. Time is indicated in minutes. Scale bar, 5 µm.

To assess which pool of cortical dynein could be generating the pulling forces for prophase centrosome positioning, we examined the localization of dynein’s cortical adaptor LIN-5^NuMA^ using an endogenous mNeonGreen (mNG) fusion (Heppert *et al*., 2018). During early prophase, LIN-5::mNG was prominently enriched on the cortex but not at apical junctions between seam cells, where co-expressed mCh::DHC-1 was the most concentrated (**Fig. 4C**). This suggests that dynein accumulates at seam cell contacts independently of LIN-5, and that this dynein pool is therefore unlikely to be involved in centrosome positioning. Instead, mCherry::DHC-1 and LIN-5::mNG were co-enriched at cortical sites away from seam cell contacts (**Fig. 4C, D**), and line scan analysis with GFP::DHC-1 and mCh::PH confirmed that cortical dynein was enriched on the seam cell side of the seam-hyp7 boundary (**Fig. 4D**), consistent with a role in cortical force generation in seam cells.

Close examination of LIN-5::mNG distribution in mitotic seam cells revealed that the anterior and posterior cortex had higher levels of LIN-5::mNG than the lateral cortex, and LIN-5::mNG tended to be displaced from the cortical region that was adjacent to separating centrosomes in early prophase (**Fig. 4E, F; S2C**). Interestingly, cortical LIN-5::mNG levels decreased as cells progressed through prophase, concomitantly with an increase of LIN-5::mNG levels at centrosomes (**Fig. S2C**). LIN-5::mNG re-appeared on the cortex during prometaphase and exhibited a prominent bipolar distribution in metaphase before its cortical levels diminished again during anaphase. The bipolar distribution of LIN-5 is consistent with the idea that pulling forces generated by LIN-5–associated dynein orient the spindle along the A–P axis, and that this cortical dynein cue already acts in early prophase to direct the migration of separating centrosomes.

To examine LIN-5 function in mitotic seam cells, we acutely inhibited LIN-5 at the larval L3 stage used the fast-acting temperature-sensitive mutant *lin-5(ev571)* (Lorson *et al*., 2000). Rapid upshift to the non-permissive temperature (26 °C) in dividing seam cells co-expressing GFP::TBB-2 and mCh::HIS-11 resulted in centrosome separation failure, while already assembled spindles became mis-oriented (**Fig. 4G**). Furthermore, LIN-5 inhibition attenuated spindle elongation in anaphase. These results support the idea that dynein contributes to spindle orientation in seam cells through its association with LIN-5 and reveals that the LIN-5–dynein pathway is required for proper spindle assembly.

Prior work demonstrated a role for the argonaute ALG-1 in seam cell spindle orientation (Wildwater *et al*., 2011), raising the possibility that cortical dynein could be regulated by pathways involving ALG-1, such as the Wnt/μ–catenin asymmetry pathway. We found that LIN-5::mNG accumulated robustly on the seam cell cortex in *alg-1(RNAi)* animals (**Fig. S2D**), which exhibited the seam defects described previously (Wildwater *et al*., 2011). This suggests that ALG-1 is not required for cortical dynein recruitment but does not rule out a role for ALG-1 in regulating dynein activity.

### GOA-1^Gαi^ is uniformly distributed on the seam cell cortex and is required for LIN-5 recruitment and prophase centrosome separation

Work in the early embryo identified the Gαi subunits GOA-1 and GPA-16 as the plasma membrane receptors for GPR-1/2, which in turn recruit LIN-5 (Colombo *et al*., 2003; Gotta *et al*., 2003; Srinivasan *et al*., 2003). To determine whether this pathway operates in seam cells, we first asked whether GOA-1 is present in seam cells using a transgene that expresses functional GOA-1, internally tagged with GFP, from *goa-1* regulatory elements (Kumar *et al*., 2021). Co-expression with mCh::HIS-11 and mCh::PH showed that GOA-1::GFP localizes uniformly to the seam cell plasma membrane at all stages of the cell cycle, which contrasts with the fluctuating levels and bipolar distribution of LIN-5::mNG (**Fig. 5A, B; Fig. S2C**). To confirm that GOA-1 acts in the dynein pathway, we fed L1 larvae with bacteria expressing dsRNA directed against *goa-1*. Subsequent examination of these larvae at the L3 stage revealed that RNAi-mediated depletion of GOA-1 in dividing seam cells de-localized LIN-5::mNG from the cell cortex, inhibited centrosome separation, and resulted in spindle mis-orientation (**Fig. 5C**). Strikingly, we found that a majority of *goa-1(RNAi)* animals ruptured upon reaching adulthood, similar to the fate of *dyci-1(11N)* animals. We conclude that GOA-1 recruits LIN-5 and dynein to the seam cell cortex and that, in contrast to the early embryo (Gotta and Ahringer, 2001), GOA-1 does not appear to function redundantly with GPA-16.

**Figure 5:**
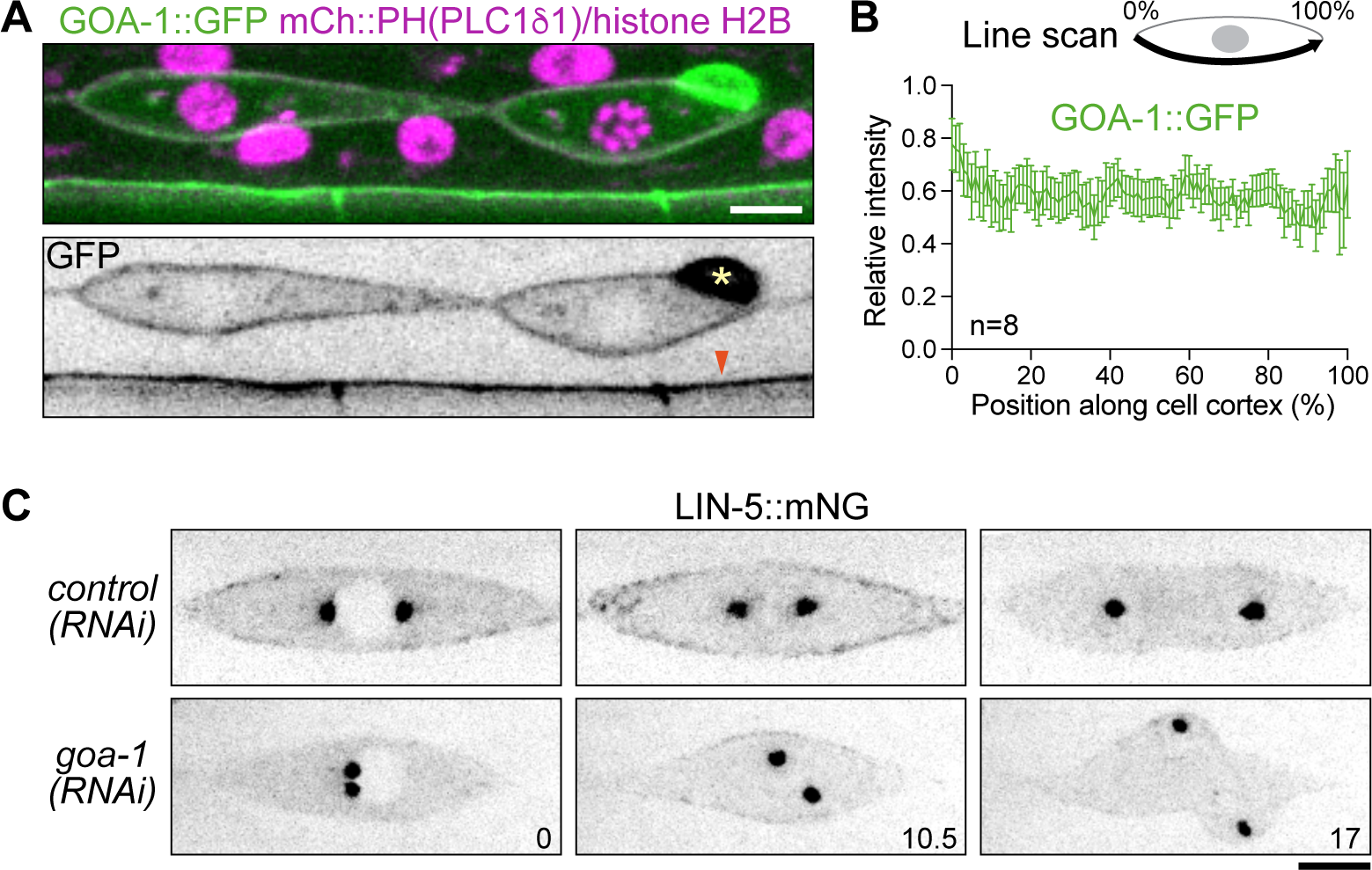
GOA-1^Gαi^ is uniformly distributed on the seam cell cortex and is required for LIN-5^NuMA^ recruitment and prophase centrosome separation. **(A)** Image of adjacent seam cells in interphase *(left)* and prophase *(right)*, co-expressing GOA-1 internally tagged with GFP, the plasma membrane marker mCh::PH(PLC181), and mCh::HIS-11 (histone H2B), showing uniform GOA-1::GFP distribution at the cell cortex. The asterisk labels a neuronal cell body adjacent to the prophase seam cell, and the arrowhead points to GOA-1::GFP signal in a neurite. Scale bar, 5 µm. **(B)** Line scan profile of GOA-1::GFP along the early prophase cell cortex, as indicated in the schematic *(top)*, in cells such as in *(A)*, showing that GOA-1::GFP distribution is uniform. Data is plotted as mean ± SEM. *n* denotes the number of cells examined (one cell per animal). **(C)** Selected images from time-lapse movies of dividing seam cells expressing LIN-5::mNG, showing that RNAi-mediated depletion of GOA-1 de-localizes LIN-5::mNG specifically from the cell cortex and impairs centrosome separation. Time is indicated in minutes. Scale bar, 5 µm.

Taken together, our results suggest that seam cells use the conserved ternary complex of GOA-1^Gαi^, GPR-1/2^LGN^, and LIN-5^NuMA^ to recruit dynein to the cell cortex in early prophase to generate pulling forces on astral microtubules that separate centrosomes and direct the path of centrosome migration along the A–P axis. The same force generation machinery subsequently maintains the spindle oriented during prometaphase and metaphase, and promotes chromosome segregation by elongating the spindle in anaphase.

### Mis-oriented divisions in the *dyci-1(ΔN)* seam result in mis-placed daughter cells and generate anucleate membrane compartments due to aberrant cytokinesis

To understand how spindle mis-orientation caused by *dyci-1(11N)* impacts the formation, positioning, and fate of daughter cells, we performed time-lapse imaging in larval L3-stage seam cells co-expressing GFP::HIS-11 and GFP::PH. Seam cell row neighbors in both control and *dyci-1(11N)* animals remained connected to each other on the apical side during mitosis and therefore maintained an elongated shape. *dyci-1(11N)* cells whose spindle was mis-oriented at anaphase onset proceeded to divide along the spindle axis, even when the resulting division axis was dramatically skewed relative to the long axis of the cell (**Fig. 6A**). Examination of cytokinesis using GFP::PH and the contractile ring marker TagRFP::ANI-1^Anillin^ revealed that the cleavage furrow in seam cells closes asymmetrically from the basal to the apical side (**Fig. 6B**). In mis-oriented *dyci-1(11N)* divisions, the cleavage plane was no longer positioned perpendicularly to the axis of tension (i.e. the seam cell row axis), which interfered with formation of a compact contractile ring (**Fig. 6C**). Occasionally, more than one region of the cortex became contractile (**Fig. 6C**), and simultaneous constriction of two cleavage furrows created apical membrane compartments between daughter cells that lacked chromosomes (**Fig. 6D**). These likely correspond to the small compartments observed with the apical junction marker GFP::DLG-1 (**Fig. 2D**). We conclude that mis-oriented divisions produce daughter cells that are positioned outside the seam cell row while promoting formation of anucleate compartments within the seam cell row.

**Figure 6:**
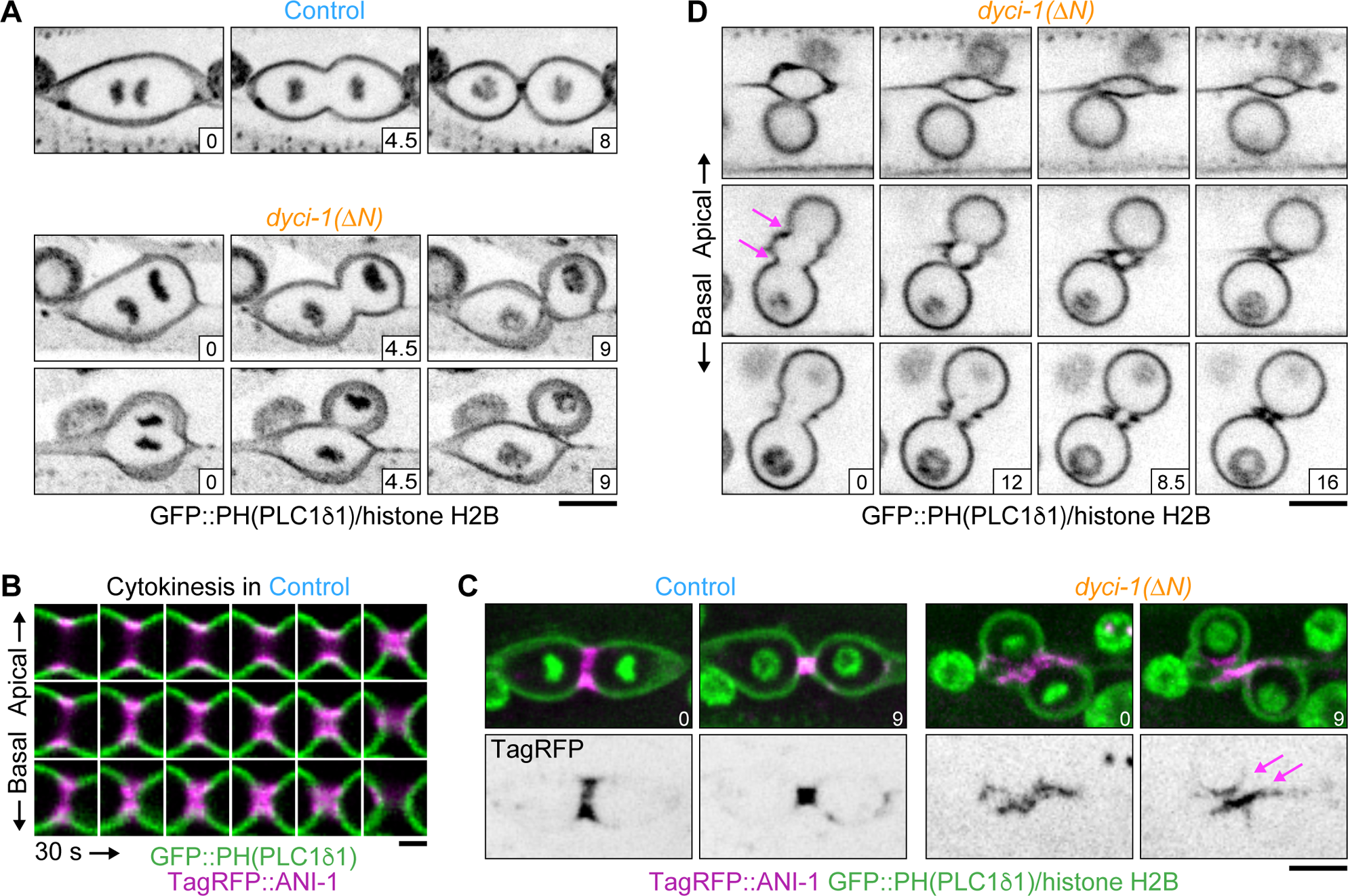
Mis-oriented divisions in the *dyci-1(ΔN)* seam result in mis-placed daughter cells and anucleate membrane compartments. **(A), (D)** Selected images from time-lapse movies of seam cells co-expressing GFP::PH(PLC181) and GFP::HIS-11 (histone H2B). Projections *(A)* or single confocal z-sections (1 µm apart) *(D)* of confocal z-stacks show mis-placement of daughter cells and formation of anucleate membrane compartments in the *dyci-1(ΔN)* mutant, respectively. Arrows in *(D)* point to two ingressing cleavage furrows in the *dyci-1(ΔN)* mutant. Time is indicated in minutes. The horizontal axis corresponds to the anterior–posterior axis in the seam. Scale bar, 5 µm. **(B)** Successive frames (30 s apart) from a time-lapse movie of cytokinesis in a seam cell co-expressing the cleavage furrow marker TagRFP::ANI-1^Anillin^, GFP::PH(PLC181), and GFP::HIS-11. Single confocal z-sections of the cleavage furrow region (0.5 µm apart) show that the furrow closes asymmetrically from the basal to the apical side. Scale bar, 2 µm. **(C)** Selected images from time-lapse movies of seam cells as in *(B)*, showing simultaneous assembly and closure of two contractile rings in the *dyci-1(ΔN)* mutant. Time is indicated in minutes. Scale bar, 5 µm.

### Mis-oriented divisions in the *dyci-1(ΔN)* seam result in incorrect inheritance of the Wnt/β**–** catenin asymmetry pathway component APR-1^APC^ and alter daughter cell fate

Following the last round of division at the L3 stage, seam cell daughters have different fates: the anterior daughter differentiates and fuses with the hyp7 cell, whereas the posterior daughter maintains seam cell identify (**Fig. 7A**). To examine how mis-oriented divisions impact cell fate, we first monitored the distribution of APR-1^APC^, a component of the Wnt/β–catenin asymmetry pathway that is asymmetrically inherited by daughter cells (Mizumoto and Sawa, 2007). In control cells, endogenous GFP-tagged APR-1 was visible on the anterior cortex by metaphase and persisted there until the end of mitosis (**Fig. 7B**). The anterior daughter therefore inherits the bulk of APR-1. In *dyci-1(11N)* cells with correctly oriented spindles, GFP::APR-1 localization was identical to that in control cells, showing that the *dyci-1(11N)* mutation does not affect APR-1 distribution *per se*. In *dyci-1(11N)* cells with mis-oriented spindles, GFP::APR-1 was still enriched on the anterior cortex at metaphase, but its distribution was skewed towards the lateral side on which the anterior spindle pole was located (**Fig. 7B**), suggesting that APR-1 distribution is influenced by the orientation of the spindle. In cases where the cleavage plane was oriented nearly parallel to the seam cell row axis, GFP::APR-1 became enriched in the ingressing cleavage furrow (**Fig. 7B**), thus ending up in both daughter cells. We conclude that mis-oriented divisions caused by *dyci-1(11N)* interfere with the correct asymmetric distribution of cell fate determinants.

**Figure 7:**
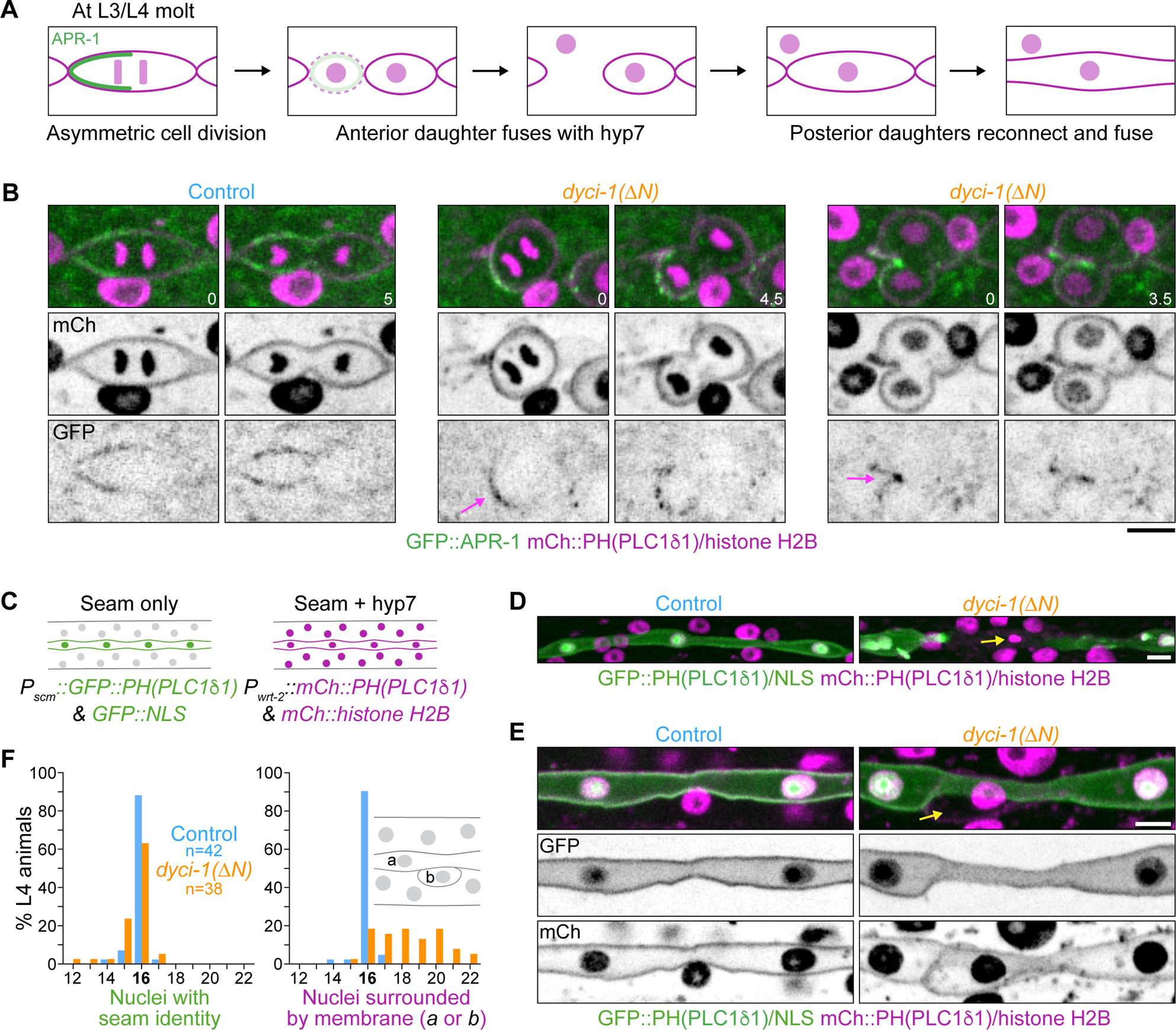
Mis-oriented divisions in the *dyci-1(ΔN)* seam result in incorrect inheritance of the Wnt/μ-catenin asymmetry pathway component APR-1^APC^ and alter daughter cell fate. **(A)** Schematic overview of asymmetric seam cell division at the L3/L4 molt and subsequent fate of anterior and posterior daughters. Selective enrichment of the Wnt/μ–catenin asymmetry pathway component GFP::APR-1 on the anterior cortex is indicated. **(B)** Selected images from time-lapse movies of seam cells co-expressing GFP::APR-1, mCh::PH(PLC181), and mCh::HIS-11 (histone H2B), showing abnormal GFP::APR-1 localization in the ingressing cleavage furrow due to spindle mis-orientation in the *dyci-1(ΔN)* mutant. Anterior is to the left. Time is indicated in minutes. Scale bar, 5 µm. **(C)** Schematic showing epidermal localization of four markers co-expressed in the cell fate reporter strain used for *(D)* – *(F)* and in Figure 8. GFP::PH(PLC181) and GFP fused to a nuclear localization signal (NLS) are expressed from the artificial *scm* promoter (*P_scm_*) to mark cells with seam identity. mCh::PH(PLC181) and mCh::HIS-11 are expressed from the *wrt-2* promoter (*P_wrt-2_*) and mark both the seam and hyp7. **(D), (E)** Images of the L4 seam in the strain described in *(C)*, showing the two types of defects encountered in the *dyci-1(ΔN)* mutant. Arrow in *(D)* points to a gap in the seam containing hyper-condensed chromatin that is characteristic of apoptosis. Arrow in *(E)* points to an individual cell adjacent to the seam that does not express the seam identity marker but is also not part of the hyp7 syncytium, indicating a problem in terminal differentiation. Scale bars, 5 µm. **(F)** Seam nuclei count in L4 animals using the markers described in (C). Control animals have 16 nuclei that express the seam identity marker and are surrounded by the plasma membrane of the seam syncytium. Most *dyci-1(ΔN)* animals have more than 16 seam–derived nuclei, and the extra nuclei correspond to individual cells such as the one shown in *(E)*. *n* denotes the number of animals examined.

To assess how mis-oriented divisions impact daughter cell fate, we constructed a strain co-expressing GFP with a nuclear localization signal (GFP::NLS) from the artificial *scm* promoter, which reports on seam cell identity (Terns *et al*., 1997), and mCh::HIS-11 from the *wrt-2* promoter, which marks all epidermal nuclei (**Fig. 7C**). The same *scm* and *wrt-2* promoters also drive expression of GFP::PH and mCh::PH from respective operons to mark the plasma membrane. In control L4 larvae, the seam syncytium contains 16 nuclei, which in our reporter strain were labeled with both GFP::NLS and mCh::HIS-11 (**Fig. 7D – F**). Furthermore, the seam was clearly delimited by GFP::PH– and mCh::PH–labelled plasma membrane. Nuclei in the surrounding hyp7 cell, by contrast, were only labeled with mCh::HIS-11. This strain therefore allowed us to identify nuclei that are physically integrated within the seam (i.e., surrounded by plasma membrane) while simultaneously evaluating seam identity of these nuclei through the *scm* reporter.

When counting nuclei positive for the *scm* reporter, we found that 32 % of *dyci-1(11N)* L4 larvae had less than 16 nuclei, compared to 10 % of control larvae (**Fig. 7F**). *dyci-1(11N)* animals with less than 16 *scm*-positive nuclei had gaps in the seam, and highly condensed mCh::HIS-11 signal was occasionally visible in these gaps (**Fig. 7D**), suggesting that gaps form in part by seam cells undergoing apoptosis. When counting the total number of seam-associated nuclei (i.e. mCh::HIS-11-positive nuclei surrounded by plasma membrane), we found that only 5 % of control L4 larvae contained more than 16 seam-associated nuclei, and in all of these cases there was only a single extra nucleus in the seam syncytium. By contrast, 79 % of *dyci-1(11N)* L4 larvae contained more than 16 nuclei, with counts ranging from 17 to 22 (**Fig. 7F**), and the extra nuclei corresponded to individual cells that were positioned outside the seam syncytium and were negative for the *scm* reporter (**Fig. 7E**). These mis-positioned cells are likely to be anterior daughters that have correctly lost expression of the seam cell identity marker but are unable to fuse with the hyp7 syncytium.

We conclude that mis-oriented division of *dyci-1(11N)* seam cells at the L3 stage results in incorrect inheritance of cell fate determinants, which in conjunction with daughter cell mis-positioning interferes with the developmental program that generates the seam syncytium with 16 nuclei at the L4 stage.

### Stretching seam cells rescues spindle mis-orientation and aberrant nuclei number in the *dyci-1(11N)* seam

A prior study demonstrated that the elongated shape of seam cells provides an important cue for correct spindle orientation (Wildwater *et al*., 2011). We therefore set out to understand how spindle mis-orientation caused by inhibition of the dynein pathway is influenced by cell shape. We used the mutations *dpy-11(e224)* and *lon-1(e185)* that produce shorter and longer animals, respectively (**Fig. 8A**), and result in corresponding changes in seam cell shape (Wildwater *et al*., 2011), which we confirmed by measuring cell length and width with the GFP::PH marker (**Fig. 8B**). This also revealed that *dyci-1(11N)* cells are rounder than controls. We then combined the *dpy-11(e224)* and *lon-1(e185)* mutations with *dyci-1(11N)* and determined the orientation of the metaphase plate, marked by GFP::HIS-11, as a proxy for spindle orientation. Spindle orientation in *dpy-11(e224)* and *lon-1(e185)* single mutants was indistinguishable from controls (**Fig. 8C**), consistent with prior work (Wildwater *et al*., 2011). Similarly, the *dpy-11(e224)*;*dyci-1(11N)* double mutant behaved the same as *dyci-1(11N)*. By contrast, combining the *lon-1(e185)* mutation with *dyci-1(11N)* resulted in significant rescue of spindle mis-orientation (**Fig. 8D**). When we counted the total number of seam-associated mCh::HIS-11-positive nuclei in L4 larvae, we found that the fraction of animals containing more than 16 nuclei was reduced from 95 % in *dyci-1(11N)* to 33 % in *lon-1(e185);dyci-1(11N)*, compared to 10 % in controls and 5 % in *lon-1(e185)* (**Fig. 8E**). We conclude that stretching seam cells in the *dyci-1(11N)* mutant to give them a more elongated mitotic shape partially suppresses spindle mis-orientation and the resulting defects in seam morphology.

**Figure 8:**
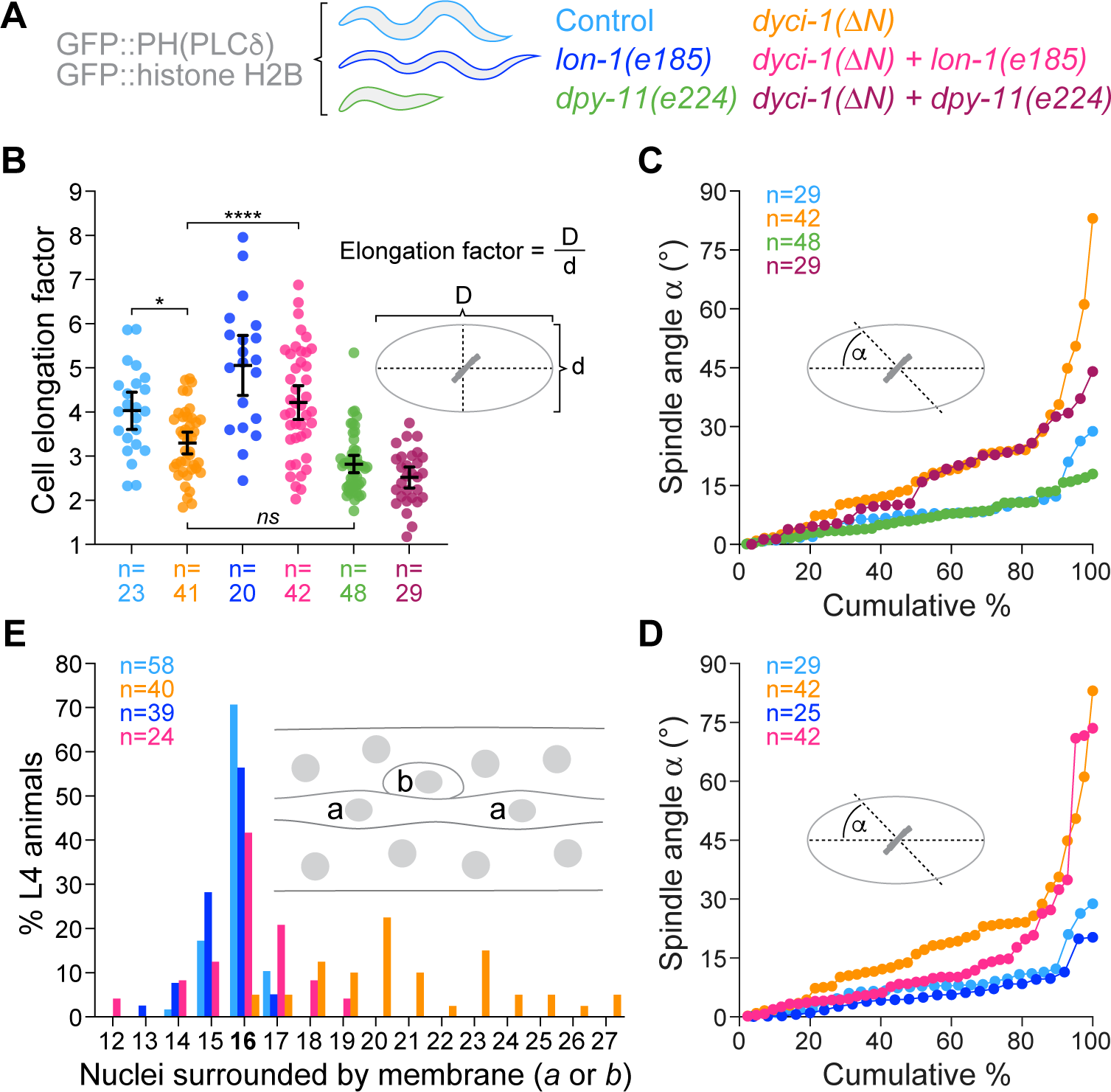
Stretching seam cells rescues spindle mis-orientation and aberrant nuclei number in the *dyci-1(11N)* seam. **(A)** Schematic illustrating the effect of the *lon-1(e185)* and *dpy-11(e224)* mutations on animal length, and the corresponding strains examined in *(B)* – *(E)*. **(B)** Quantification of the seam cell elongation factor at metaphase, calculated as shown in the schematic. Data points correspond to measurements in individual cells and bars show the mean ± 95% CI. *n* denotes the total number of cells examined in at least 10 animals. Statistical significance was determined by ordinary one-way ANOVA followed by Šídák’s multiple comparison test. *\*\*\*\*P* < 0.0001; *\*P* < 0.05; *ns* = not significant, *P* > 0.05. **(C), (D)** Quantification of seam cell spindle orientation relative to the anterior–posterior axis at metaphase, showing that although both *dyci-1(ΔN)* and *dpy-11(e224)* increase cell roundness, only *dyci-1(ΔN)* causes spindle mis-orientation, and that stretching cells with *lon-1(e185)* partially rescues spindle mis-orientation in the *dyci-1(ΔN)* background. Data points correspond to measurement in individual cells. *n* denotes the total number of cells examined in at least 10 animals. **(E)** Seam nuclei count in L4 animals using the markers described in Figure 7C and the criteria outlined in the schematic, showing that *lon-1(e185)* reduces the number of extra nuclei in the *dyci-1(ΔN)* background. *n* denotes the number of animals examined.

## DISCUSSION

The *C. elegans* epidermis is a model to study stem cell-like divisions in a simple epithelium. Here we characterized the role of the microtubule motor dynein during the last round of asymmetric seam cell divisions that give rise to the seam syncytium. Our results show that dynein acts together with elongated cell shape to ensure spindle assembly occurs along the A–P axis, and we demonstrate that this is crucial for developing a functional seam. We further show that dynein’s contribution is mediated by the conserved cortical Gαi–LGN–NuMA pathway and begins in prophase, when the trajectory followed by separating centrosomes establishes the orientation of the spindle.

### The importance of oriented division for seam development

A previously characterized mutant of the argonaut *alg-1* was found to alter the seam cell division axis and to produce a dis-organized seam, but this mutant also perturbed multiple other processes that are important for correct development of the epidermis, including the timing of seam cell divisions and the Wnt/β–catenin asymmetry pathway (Wildwater *et al*., 2011). Partial inhibition of dynein, the most downstream component of the cortical force generation machinery, enabled us to more directly address the importance of oriented division for seam cell development. The hypomorphic dynein mutant *dyci-1(11N)* is mild enough to support spindle assembly and error-free chromosome segregation of seam cells at the L3 stage but penetrant enough to produce significant spindle mis-orientation. The immediate consequences of spindle mis-orientation are two–fold: daughter cells are placed outside the seam cell row, and anucleate compartments form due to mis-positioning of cytokinetic furrows, which are forced to close against the axis of tension.

How are the defects caused by spindle mis-orientation at the L3 stage related to the subsequent defects observed in the mature seam syncytium at the L4 stage? A prominent defect in the mature *dyci-1(11N)* seam is the presence of seam–derived cells that persist adjacent to the seam syncytium but are negative for a seam identity marker. These cells are most likely anterior daughters that are unable to fully differentiate, which involves fusion with the hyp7 syncytium. This suggests that mis-oriented division alters seam cell fate, consistent with the observation that the Wnt/β–catenin asymmetry pathway component APR-1^APC^ becomes inappropriately distributed to daughter cells following mis-oriented division. Support for a causal relationship between mis-oriented division and failure to fuse with hyp7 is provided by our observation that stretching *dyci-1(11N)* cells with the *lon-1(e185)* mutant not only rescues spindle mis-orientation in L3 larvae but also reduces the number of the extra seam-derived cells at the L4 stage.

The other major defects observed in the mature *dyci-1(11N)* seam are anucleate compartments and large gaps, which is predicted to be particularly detrimental since the seam secretes components that make up the cuticle to ensure impermeability of the epidermis (Chisholm and Xu, 2012). Gaps may form when anucleate compartments produced by mis-oriented divisions degenerate, or when mis-placed posterior daughter cells are unable to re-connect after anterior daughters have fused with hyp7. Taken together, our results show that seam development is highly sensitive to mis-oriented divisions, highlighting the importance of redundant mechanisms for spindle orientation, as discussed below.

### Dynein’s contribution to spindle orientation in the context of cell shape and Wnt signalling

Hertwig’s rule states that spindles orient along the long axis of the cell, and a previous study established the importance of elongated cell shape for oriented seam cell division in the context of perturbed Wnt signalling (Wildwater *et al*., 2011). Specifically, inhibition of the argonaut ALG-1 perturbed Wnt signalling and made cells rounder, and the resulting mis-oriented divisions could be rescued by stretching cells with the *lon-1(e185)* mutation. Furthermore, combining the *dpy-11(e224)* mutation, which makes seam cells rounder, with inhibition of Wnt signalling also resulted in mis-oriented divisions (Wildwater *et al*., 2011). Our findings are analogous in that the *dyci-1(11N)* mutation produces L3 seam cells that are rounder than controls, and stretching *dyci-1(11N)* cells with the *lon-1(e185)* mutation partially rescues spindle mis-orientation. The *dpy-11(e224)* mutant also shows that roundness *per se* is not sufficient to induce spindle mis-orientation, in agreement with prior results (Wildwater *et al*., 2011). Taken together, this suggests that spindle mis-orientation in *dyci-1(11N)* cells is a combined consequence of dynein inhibition and increased cell roundness. This re-enforces the notion that redundant geometric and cortical cues exist to provide robust control over the cell division axis (Wildwater *et al*., 2011).

Since both dynein (this study) and Wnt signaling (Wildwater *et al*., 2011) contribute to spindle orientation in seam cells, the question arises whether Wnt signalling regulates cortical dynein. Wnt signaling has been shown to direct dynein to the cortex in *D. melanogaster* and vertebrate cells, albeit in an LGN-independent manner (Lechler and Mapelli, 2021). We did not observe an obvious reduction in cortical LIN-5^NuMA^ levels after *alg-1(RNAi)*, which perturbs Wnt signaling (Wildwater *et al*., 2011), and Wnt signalling in the four-cell embryo, while required for spindle orientation, was shown to be dispensable for LIN-5 enrichment at the P2–EMS cell boundary (Srinivasan *et al*., 2003; Heppert *et al*., 2018). Nevertheless, a more detailed examination is needed to clarify whether Wnt signaling contributes to spindle orientation in seam cells through the cortical dynein pathway or in a dynein-independent manner, for example through APR-1–mediated microtubule stabilization (Sugioka *et al*., 2011).

### Dynein–dependent cortical force generation separates prophase centrosomes in seam cells

We find that the depletion of GOA-1 ^Gαi^ by RNAi has a strong inhibitory effect on prophase centrosome separation, suggesting that in seam cells GOA-1 function is not redundant with that of its paralog GPA-16. This contrasts with the situation in the one-cell embryo, where co-depletion of GOA-1 and GPA-16 slows but does not prevent prophase centrosome separation (De Simone *et al*., 2016). In the one-cell embryo, cortex–associated dynein acts together with nucleus-associated dynein to separate centrosomes (De Simone *et al*., 2016), whereas in seam cells we did not detect enrichment of dynein at the nuclear envelope. The apparent lack of nucleus-associated dynein may explain why the cortical dynein pathway in seam cells plays a more prominent role in centrosome separation compared to the one-cell embryo.

#### Cortical dynein in seam cells directs prophase centrosome migration along the A–P axis

At the end of mitosis, the duplicated centriole pair in each daughter cell naturally comes to rest on the axis of division. Consequently, if both centrosomes migrate evenly to opposite sides of the nucleus during the next prophase, the resulting centrosome–centrosome axis becomes positioned orthogonally to the prior axis of division. In the blastomeres of the early embryo, centrosomes do indeed migrate evenly to opposite sides of the nucleus, which gives rise to the orthogonal division pattern of AB cells (Hyman and White, 1987). The P0 blastomere and its descendants (P1, P2, and EMS), however, need to divide along the A–P axis. In these cells, the centrosome–nucleus complex is rotated into the correct position once centrosomes have separated (Hyman and White, 1987). We find that seam cells use a different strategy to ensure successive divisions along the A–P axis. Instead of rotating the centrosome–nucleus complex, the migration path of separating centrosomes is directed along the A–P axis, which typically means that one of the centrosomes stays in place while the other moves to the opposite side of the nucleus. Our analysis of the *dyci-1(11N)* mutant shows that dynein plays an important role in this process. We speculate that directed centrosome migration along the A–P axis is facilitated by the bipolar enrichment of the cortical dynein adaptor LIN-5^NuMA^, which is discussed below.

### Localization dynamics of the dynein force generation machinery at the seam cell cortex

To the best of our knowledge, Gαi localization in mitosis has not been examined by live cell imaging in any organism. We took advantage of a recently generated functional GFP-tagged version of GOA-1 (Kumar *et al*., 2021) to examine GOA-1 localization in seam cells. This revealed that GOA-1 is uniformly distributed on the cortex throughout mitosis, which contrasts with the fluctuating levels and uneven distribution of LIN-5^NuMA^. Similar observations were made by immunofluorescence in chick neuroepithelial and HeLa cells (Kiyomitsu and Cheeseman, 2012; Peyre *et al*., 2011), in which Gαi, but not LGN–NuMA, is uniformly distributed on the metaphase cortex. Since we can only observe the total GOA-1 pool, it is possible that GOA-1–GDP, the specific GOA-1 pool that interacts with GPR-1/2^LGN^ (Srinivasan *et al*., 2003; Gotta *et al*., 2003; Colombo *et al*., 2003), exhibits a distribution similar to that of LIN-5. Another possibility is that GOA-1–GDP is uniformly distributed and the interaction between GOA-1–GDP and GPR-1/2 (for which we had no live imaging probe), or the interaction between GPR-1/2 and LIN-5, is subject to regulation.

Work in HeLa cells showed that the RanGTP gradient generated by mitotic chromosomes excludes cortical LGN–NuMA from regions near the spindle midzone (Kiyomitsu and Cheeseman, 2012). A chromosomal RanGTP gradient in seam cells could explain the prominent bipolar LIN-5 distribution on the metaphase and early anaphase cortex but not its cortical distribution prior to NEBD. In prophase, we find that LIN-5 tends to be depleted from cortical regions that are closest to the separating centrosomes and that LIN-5 is preferentially concentrated in the polar regions of the elongated seam cell, particularly on the side of the nucleus opposite to the centrosome pair. We speculate that cell pole–enriched localization of LIN-5–dynein directs centrosome migration along the A–P axis. In HeLa cells, the proximity of spindle poles to the cortex displaces cortical dynein, albeit downstream of NuMA, through the pole–localized kinase Plk1 (Kiyomitsu and Cheeseman, 2012). It would therefore be interesting to examine whether PLK-1 plays a role in displacing LIN-5–dynein from the centrosome–proximal seam cell cortex during centrosome migration. Interestingly, the progressive increase of LIN-5 levels at prophase centrosomes is accompanied by a corresponding reduction of cortical LIN-5 levels, suggesting that LIN-5–dynein may remove itself from the prophase cortex through transport along astral microtubules. Removal from the cortex along astral microtubules was previously documented for GPR-1/2 in the P2 cell of the four–cell embryo, although in this case removal occurs only once spindle orientation has completed and serves to prevent hyperaccumulation of GPR-1/2 (Werts *et al*., 2011).

### Additional roles of dynein in seam cells beyond cell division

Although spindle mis-orientation is significantly rescued by stretching *dyci-1(11N)* seam cells, *dyci-1(11N);lon-1(e185)* animals still die prematurely by epidermal rupture and still exhibit gaps in the seam (data not shown). While this may in part reflect incomplete rescue of spindle mis-orientation by cell stretching, it may also indicate that dynein has additional important roles in the epidermis. For example, we find that dynein, but not its cortical adaptor LIN-5, accumulates prominently at apical junctions that connect neighboring seam cells, which is where microtubule minus-ends concentrate during interphase (Wang *et al*., 2015). We also find that several cargo adaptors for dynein involved in membrane trafficking accumulate at apical junctions along with the motor (data not shown). It is therefore conceivable that dynein–driven membrane trafficking directed to apical junctions is required for maintenance of proper cell–cell adhesion within the seam cell row, which is under considerable tension. Dynein–driven membrane trafficking may also be required for proper secretion of cuticle components. Consistent with this possibility, a null mutant of the small GTPase RAB-6.2, which marks golgi-derived exocytotic vesicles that have been identified as dynein cargo in other species, exhibits cuticle integrity defects (Kim *et al*., 2019). It will be interesting to investigate these potential contributions of dynein to epidermal development in future work.

## MATERIALS & METHODS

### C. elegans

Strains (**Supplemental Table 1**) were maintained at 20 °C, except for *lin-5(ev571)* (16 °C), on standard nematode growth media (NGM) plates seeded with OP50 bacteria. The *dyci-1* N-terminal deletion mutant *prt152* was generated by CRISPR/Cas9-mediated genome editing as described (Arribere *et al*., 2014; Paix *et al*., 2014), using two crRNAs targeting the genomic regions 5’ TTTCAGAATGTCAGAACTGAGG 3’ and 5’ CCGTTCCACAAATGAAAATGGA 3’ and a single-stranded oligonucleotide repair template. The 99-bp deletion was confirmed by sequencing. To remove potential off-target mutations, *dyci-1(11N)* animals were outcrossed 6x with the wild-type N2 strain. Other modifications were subsequently introduced by mating. The *dyci-1(11N)* mutation was maintained in a heterozygous state using the GFP-marked balancer nT1 [qIs51]. Homozygous F1 progeny from balanced heterozygous mothers was identified by the absence of GFP fluorescence in the pharynx. A Mos1 transposon-based strategy (Frøkjaer-Jensen *et al*., 2012) was used to generate prtSi193, prtSi199, prtSi222, prtSi239, and prtSi262. Transgenes were cloned into pCFJ151 for insertion on chromosome II (ttTi5605 locus) or chromosome V (oxTi365), and transgene integration was confirmed by PCR.

### Synchronization at the L1 stage

Adult animals were washed once with M9 buffer, once with 0.1 M NaCl, bleaching solution (67 % 0.1 M NaCl, 22 % house-hold bleach, 11 % 5 N NaOH) was added, and the worm suspension was vortexed for 10 min. Embryos were pelleted, washed 4 x with M9 buffer, and allowed to develop and hatch in M9 buffer overnight at room temperature. For release from L1 arrest, animals were transferred to an NGM plate with bacteria.

### Life span

Synchronized animals were collected 24 h after release from L1 arrest and transferred every 2 days to a new NGM plate with bacteria. Animals were examined every day and scored as dead if they did not respond to touch and if there was no evidence of pharyngeal pumping. Animals that escaped or were found dead on the edge of the plate were excluded from the assay.

### Permeability

Synchronized animals at the adulthood day 1 stage (72 h after release from L1 arrest) were incubated in a depression slide well with 1 µg/mL of the dye Hoechst 33342 (Invitrogen) for 15 min in the dark. Animals were washed twice with 0.7x Egg Salts buffer (1x Egg Salts buffer is 5 mM HEPES pH 7.4, 118 mM NaCl, 40 mM KCl, 3.4 mM MgCl_2_, 3.4 mM CaCl_2_), immobilized with 5 mM levamisole for 10 min, and mounted on a freshly prepared 2 % (w/v) agarose pad for imaging. Animals were scored as permeable if at least three epidermal nuclei were stained with Hoechst 33342.

### Animal length

48 h after release from L1 arrest, animals were transferred to a new NGM plate with a thin layer of bacteria. Images were taken at 20 °C using an SMZ 745T stereoscope (Nikon) equipped with a QIClic CCD camera (QImaging) and controlled by Micro-Manager software (Open Imaging). Animal length was determined by tracing a segmented line along the body using Fiji software.

### RNA interference

RNAi was performed by feeding animals with HT115 *E. coli* bacteria expressing dsRNA. The L4440 plasmid containing part of the *goa-1* genomic sequence was obtained from the Ahringer library (Kamath and Ahringer, 2003; distributed by Source BioScience). L4440 containing the *alg-1* sequence was built by inserting an *alg*-1 cDNA fragment amplified with primers 5’ TCCATGCTTCTGCAAGTACG 3’ and 5’ ACCTGCACAGCTCTAGCCAT 3’ into the EcoRV restriction site. To prepare feeding plates, bacteria were grown in LB containing 12.5 μg/mL tetracycline and 100 μg/mL ampicillin until an OD_600_ of 1.6, then pelleted at 2500 x g for 10 min and resuspended in LB with 12.5 μg/ml tetracycline, 100 μg/mL ampicillin, and 1 mM IPTG. Resuspended bacteria were spread onto NGM agar plates (∼75 μL/plate) previously soaked with 100 μL of a 1:1:1 mixture of 100 mg/mL ampicillin, 5 mg/mL tetracycline and 1 M IPTG. Plates were dried in the dark at room temperature for 1 – 3 days to induce RNA expression. L1 animals were placed on the plates and incubated at 20 °C until imaging.

### Immunoblotting

200 adults were collected into cold 0.7x Egg Salts buffer, washed 3 times with cold 0.7x Egg Salts buffer, and washed once with 0.7x Egg Salts buffer containing 0.05 % Triton X-100. To 100 µL of worm suspension, 33 µL 4x SDS-PAGE sample buffer [250 mM Tris-HCl pH 6.8, 30 % (v/v) glycerol, 8 % (w/v) SDS, 200 mM DTT and 0.04 % (w/v) bromophenol blue] and 20 μL glass beads were added. Samples were incubated for 3 min at 95 °C and vortexed for 2 x 5 min. Supernatants were collected after centrifugation at 20000 x g for 1 min at room temperature. Proteins were resolved by SDS-PAGE and transferred to 0.2 μm nitrocellulose membranes (Hybond ECL, Amersham Pharmacia Biotech). Membranes were rinsed 3 x with TBS (50 mM Tris-HCl pH 7.6, 145 mM NaCl), blocked with 5 % non-fat dry milk in TBST (TBS with 0.1 % Tween-20) and probed at 4 °C overnight with the following primary antibodies: mouse monoclonal anti-FLAG M2 (Sigma-Aldrich F3165, 1:1000), mouse monoclonal anti-α-tubulin B512 (Sigma-Aldrich T5168, 1:5000), rabbit polyclonal anti-DYCI-1 GC1 (Barbosa *et al*., 2017, 1:2000), rabbit polyclonal anti-DHC-1 GC4 (Simões *et al*., 2018; 1:1000), and rabbit polyclonal anti-DLI-1 GC11 (made in-house, 1:2500). Membranes were washed 3 x with TBST, incubated with goat polyclonal anti-mouse IgG (Jackson ImmunoResearch 115-035-044, 1:10000) or anti-rabbit IgG (Jackson ImmunoResearch 111-035-003, 1:10000), coupled to HRP, for 1 hour at room temperature, and washed again 3 x with TBST. Proteins were detected by chemiluminescence using Pierce ECL Western Blotting Substrate (Thermo Scientific) and X-ray film (Fuji).

### DLI-1 antibody

Recombinant GST::DLI-1 (residues 1-368) was purified from bacteria and injected into rabbits (GeneCust). Antibodies were affinity purified against DLI-1(1-368) coupled to a 1-mL HiTrap *N*-hydroxysuccinimide column (GE Healthcare) according to the manufacturer’s instructions.

### Microscopy

Animals were immobilized with 5 mM levamisole for 10 min, mounted on a freshly prepared 2 % (w/v) agarose pad, and covered with an 18 mm × 18 mm coverslip (No. 1.5H, Marienfeld). For time-lapse imaging, the coverslip was sealed with VALAP (1:1:1 Vaseline: lanolin: paraffin) to prevent desiccation. Imaging was performed at 20 °C with two microscopes: a Zeiss Axio Observer, equipped with an Orca Flash 4.0 camera (Hamamatsu) and a Colibri 2 light source (Zeiss), controlled by ZEN software (Zeiss); and a Nikon Eclipse Ti coupled to an Andor Revolution XD spinning disk confocal system, composed of an iXon Ultra 897 CCD camera (Andor Technology), a solid-state laser combiner (ALC-UVP 350i, Andor Technology), and a CSU-X1 confocal scanner (Yokogawa Electric Corporation), controlled by Andor IQ3 software (Andor Technology).

#### Seam morphology, cell fate markers, cytokinesis, and epidermal microtubules

For quantification of seam morphology, late-L4 stage DLG-1::GFP animals were imaged on the Axio Observer microscope with a 40x NA 1.3 Plan-Neofluar objective. For imaging of GFP::NLS/GFP::PH/mCh::HIS-11/mCh::PH and TagRFP::ANI-1/GFP::PH/GFP::H2B, 5-µm z-stacks (step size 0.1 – 0.5 µm) were acquired on the spinning disk confocal system using a 60x NA 1.4 Plan-Apochromat objective. For imaging of GFP::TBB-2, z-stacks (step size 0.1 µm) were acquired on the spinning disk confocal system using a 100x NA 1.45 Plan-Apochromat objective.

#### Live imaging of dividing seam cells

Developmentally synchronized animals were imaged 30 h after release from L1 arrest. Since feeding with HT115 bacteria slightly delays development, animals in RNAi experiments were imaged 36 h after release from L1 arrest. Imaging was performed on the spinning disk confocal system using a 60x NA 1.4 Plan-Apochromat or 100x NA 1.45 Plan-Apochromat objective. A 3- to 5-µm z-stack was acquired every 30 s from the start of centrosome separation until cytokinesis.

#### CherryTemp

Rapid temperature shifts were performed using the CherryTemp heater-cooler (Cherry Biotech) coupled to the spinning disk confocal system. Animals were kept at 16 °C for 55 h after release from L1 arrest and mounted on a thin 2 % (w/v) agarose pad, which was inverted onto another pad and cropped to fit the 20-µm spacer. This setup was transferred to the CherryTemp chip, engravements facing down, pad centred on the thermalising pattern. Temperature was initially set to 16 °C (permissive temperature). After identification of dividing cells, temperature was upshifted to 26 °C (restrictive temperature), the focus was adjusted, and live imaging was performed with a 60x NA 1.4 Plan-Apochromat objective.

### Image analysis

Image analysis was performed with Fiji and Imaris (Oxford Instruments) software.

#### Microtubule bundle density

After maximum intensity projection of the z-stack, the ‘Plot Profile’ function in Fiji was used on a free-hand line of approximately 40 µm drawn along the A–P axis in the dorsal or ventral region. Microtubule bundle density was calculated by dividing the number of distinct GFP::TBB-2 peaks by the length of the line.

#### Spindle orientation and length

In GFP::TBB-2/mCh::HIS-11 movies, the XYZ coordinates of centrosomes and spindle poles were determined automatically using Imaris software (Oxford Instruments), and the tracks were manually verified. In GFP::PH/GFP::HIS-11 movies, the XY coordinates of the two outermost chromosomes of the metaphase plate were determined using the MTrackJ plugin in Fiji to define the metaphase plate axis. The A–P axis was defined by the coordinates of two points on the seam cell long axis.

#### Cell shape

The elongation factor was determined by dividing the length of the cell by its width. Measurements were performed in the frame before anaphase onset in an apical z-plane where the junctions that connect seam cells are visible.

#### Cortical line profiles

For line profiles across the hyp7–seam cell border, a straight line was drawn perpendicularly to the border in unprojected z-stacks, starting in the hyp7 cell. The line was 4 µm long and 10 pixels wide, except for mCh::DHC-1/LIN-5::mNG (3 µm long and 4 pixels wide). The line was then centered on the border, and the fluorescence profile was recorded using the ‘Plot Profile’ function in Fiji. The average of the first 5 signal intensity values was designated as background and was subtracted from the line profile. GFP/mCherry profiles were aligned relative to the peak mCherry signal, profiles of replicates were averaged, normalized to the maximum average, and plotted as mean ± SEM. GFP::DHC-1 values in *dyci-1(11N)* were normalized to the maximum average of the control profile to allow for relative comparison of intensities.

For line profiles along the hyp7–seam cell border, a segmented line was drawn along the lateral side of the cell in 2 - 3 successive z-planes for a total of 4 - 6 line profiles per cell. The line was 4 pixels and 5 pixels wide for LIN-5::mNG and GOA-1::GFP, respectively. Signal intensity profile was normalized to line length and averaged over 1 % intervals. For LIN-5::mNG, the lowest value in the profile plot was considered background. For GOA-1::GFP, the mean signal intensity in a box in the seam cell cytoplasm served as background. After background subtraction, all profiles from an individual cell were averaged, and resulting profiles from different cells were plotted as mean ± SEM.

### Statistical analysis

Statistical analysis was performed with Prism 9.0 software (GraphPad). Statistical significance was determined by ordinary one-way ANOVA followed by Šídák’s multiple comparison test, or by a two-sided Mann-Whitney test, where \*\*\*\**P* < 0.0001, \*\*\**P* < 0.001, \*\**P* < 0.01, \**P* < 0.05, and *ns* = not significant, *P* > 0.05. The analytical method is specified in the figure legends.

## ACKNOWLEDGEMENTS

The authors thank Eurico Morais de Sá (i3S Porto) for critical reading of the manuscript; Michael Koelle (Yale University School of Medicine) and Karen Oegema (University of California San Diego) sharing strains; and WormBase for providing data and tools. Some strains were provided by the *Caenorhabditis* Genetics Center (CGC), which is funded by the NIH Office of Research Infrastructure Programs (P40 OD010440). This project was funded by the European Research Council under the European Union’s Seventh Framework Programme (ERC-2013-StG-338410-DYNEINOME) and by the Fundação para a Ciência e a Tecnologia (FCT)/Ministério da Ciência, Tecnologia e Ensino Superior (PTDC/BIA-CEL/1321/2021). R. G. and A. C. are supported by FCT Principal Investigator positions (CEECIND/00333/2017 and CEECIND/01967/2017, respectively), D. J. B. is supported by an FCT Junior Researcher position (DL57/2016/CP1355/CT0001), and C. C. is supported by an FCT PhD fellowship (SFRH/BD/144877/2019). The authors declare no competing financial interests.

---

**Figure S1:**
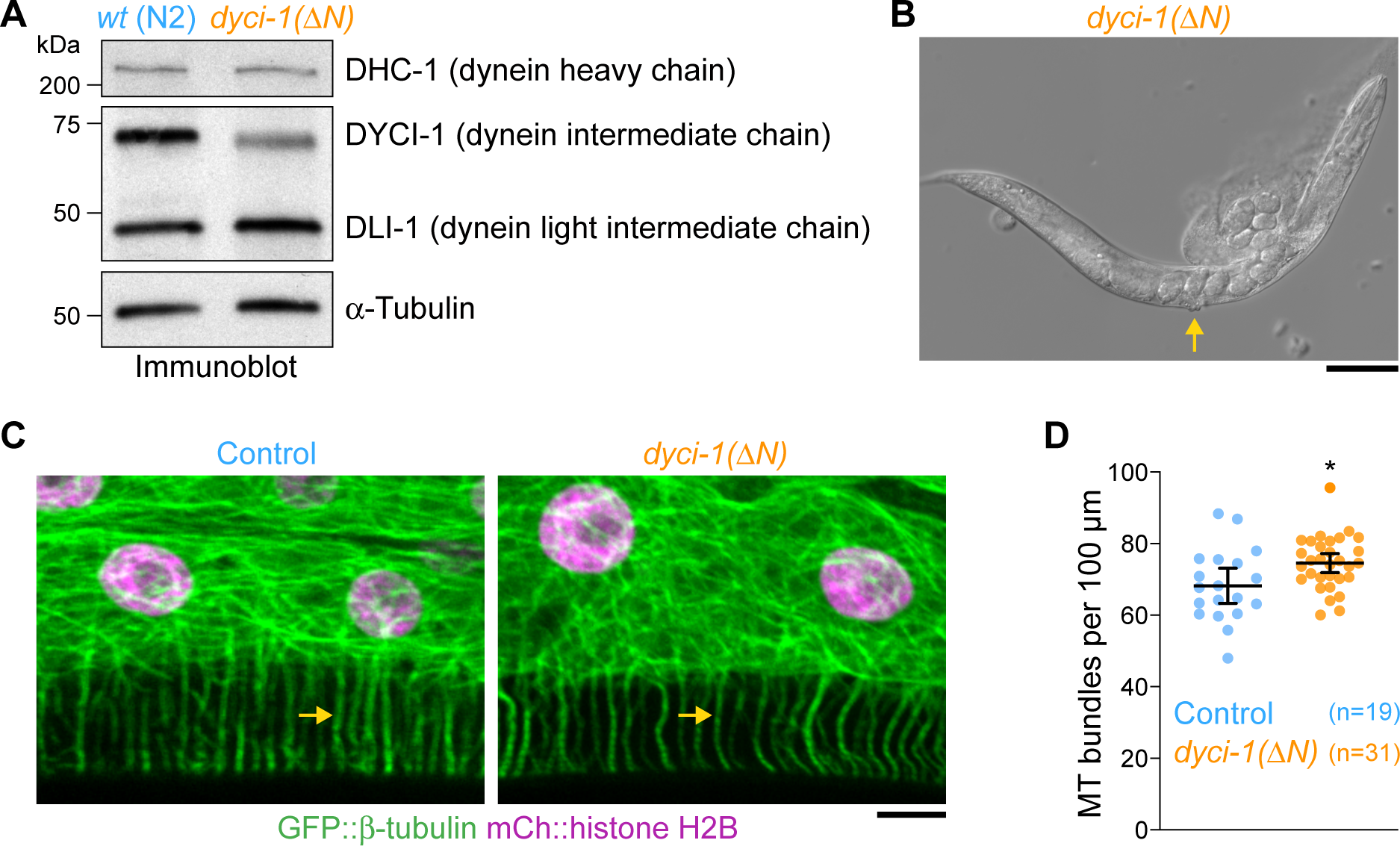
Extra data related to the characterization of the *dyci-1(ΔN)* mutant in Figures 1 and 2. **(A)** Immunoblot of adult animals, showing that *dyci-1(ΔN)* reduces the levels of DYCI-1 but not other dynein subunits. Anti–α-tubulin antibody serves as the loading control. Molecular weight is indicated in kilodaltons (kDa). **(B)** Differential interference contrast image of a ruptured adult *dyci-1(ΔN)* animal with embryos, showing that rupturing did not occur through the vulva, which remains intact (arrow). Scale bar, 100 µm. **(C)** Images of hyp7 at the L4 stage in animals co-expressing GFP::TBB-2 (β-tubulin) and mCh::HIS-11 (histone H2B), showing correctly organized microtubule bundles (arrow) in the *dyci-1(ΔN)* mutant. Scale bar, 5 µm. **(D)** Quantification of microtubule bundle density in hyp7, using images such as shown in *(C)*. Data points correspond to counts in individual animals with bars showing the mean ± 95% CI. *n* denotes the number of animals examined. Statistical significance was determined by the Mann-Whitney test. *\*P* < 0.05.

**Figure S2:**
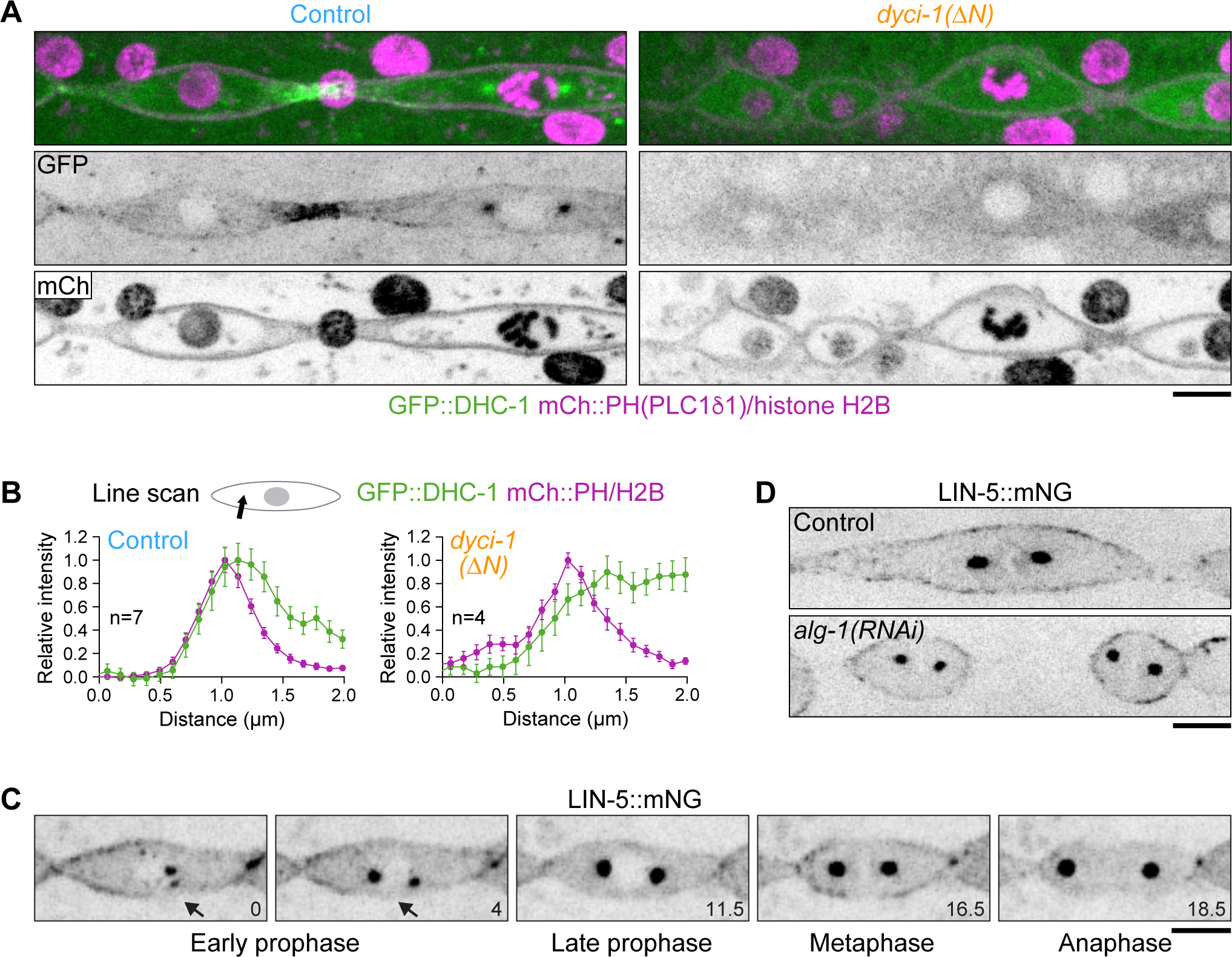
Extra data related to the characterization of seam cell dynein in Figures 4 and 5. **(A)** Images of an interphase *(left)* and late prophase *(right)* seam cell co-expressing GFP::DHC-1 (dynein heavy chain), the plasma membrane marker mCh::PH(PLC1δ1), and mCh::HIS-11 (histone H2B), showing de-localization of GFP::DHC-1 from centrosomes, the cell cortex, and cell–cell junctions in the *dyci-1(ΔN)* mutant. Scale bar, 5 µm. **(B)** Line scan profiles across the prophase seam cell cortex, as illustrated in the schematic, measured in images such as in *(A)*, showing increased cytosolic levels of GFP::DHC-1 due to de-localization from sub-cellular sites. Data is plotted as mean ± SEM. *n* denotes the total number of cells examined in at least 4 animals. **(C)** Selected images from a time-lapse movie of a dividing seam cell expressing LIN-5::mNG, showing progressively decreasing cortical levels in prophase (time points 0, 4, and 11.5 minutes), robust bipolar distribution on the cortex in metaphase (time point 16.5 minutes), and decreased cortical levels in anaphase (time point 18.5 minutes). Arrows point to the absence of cortical LIN-5::mNG near prophase centrosomes. Scale bar, 5 µm. **(D)** Images of dividing seam cells expressing LIN-5::mNG, showing that LIN-5::mNG remains on the cortex after *alg-1(RNAi)*, which results in a disorganized seam, due in part to perturbed Wnt signaling and spindle mis-orientation (Wildwater *et al*., 2011). Scale bar, 5 µm.

**Table S1:** *C. elegans* strains. Strain Genotype.

| **Strain** | **Genotype** |
| --- | --- |
| **N2** | N2 Bristol wild-type isolate |
| **OD1741** | *ItSi570[Pdpy-7::GFP::tbb-2::mCherry::his-11; cb-unc-119(+)] I; unc-119(ed3) III* |
| **OD961** | *ItSi249[Pdlg-1delta7::dlg-1::GFP; cb-unc-119(+)] I; unc-119(ed3) III* |
| **LP585** | *lin-5(cp288[lin-5::mNG-C1^3xflag]) II* |
| **ZAN370** | *ani-1(syb4317[TagRFP::ani-1]) III* |
| **GCP2** | *dyci-1(tm4732) IV/nT1[qIs51] (IV;V)* |
| **GCP786** | *dyci-1(prt152[Δ4-36]) IV/nT1[qIs51] (IV;V)* |
| **GCP893** | *ItSi249[Pdlg-1delta7::dlg-1::GFP; cb-unc-119(+)] I; unc-119(ed3) III; dyci-1(prt152[Δ4-36]) IV/nT1[qIs51] (IV;V)* |
| **GCP902** | *ItSi570[Pdpy-7::GFP::tbb-2::mCherry::his-11; cb-unc-119(+)] I; unc-119(ed3) III; dyci-1(prt152[Δ4-36]) IV/nT1[qIs51] (IV;V)* |
| **GCP972** | *prtSi193[Pdpy-7::dyci-1(wt)::3xFLAG; cb-unc-119(+)] II; unc-119(ed3) III; dyci-1(prt152[Δ4-36]) IV/nT1[qIs51] (IV;V)* |
| **GCP984** | *prtSi193[Pdpy-7::dyci-1(Δ4-36)::3xFLAG; cb-unc-119(+)] II; unc-119(ed3) III; dyci-1(prt152[Δ4-36]) IV/nT1[qIs51] (IV;V)* |
| **GCP1055** | *prtSi222[Pwrt-2::GFP::his-58::GFP::PH(PLC1δ1); cb-unc-119(+)] II; unc-119(ed3) III* |
| **GCP1062** | *prtSi222[Pwrt-2::GFP::his-58::GFP::PH(PLC1δ1); cb-unc-119(+)] II; unc-119(ed3) III; dyci-1(prt152[Δ4-36]) IV/nT1[qIs51] (IV;V)* |
| **GCP1137** | *dhc-1(he264[N-terminal GFP]) I; prtSi239[Pwrt-2::mCherry::his-58::mCherry::PH(PLC1δ1); cb-unc-119(+)] II; unc-119(ed3) III* |
| **GCP1159** | *dhc-1(he264[N-terminal GFP]) I; prtSi239[Pwrt-2::mCherry::his-58::mCherry::PH(PLC1δ1); cb-unc-119(+)] II; unc-119(ed3) III; dyci-1(prt152[Δ4-36]) IV/nT1[qIs51] (IV;V)* |
| **GCP1171** | *ItSi249[PdIg-1delta7::dlg-1::GFP; cb-unc-119(+)] I; prtSi239[Pwrt-2::mCherry::his-58::mCherry::PH(PLC1δ1); cb-unc-119(+)] II; unc-119(ed3) III* |
| **GCP1174** | *prtSi222[Pwrt-2::GFP::his-58::GFP::PH(PLC1δ1); cb-unc-119(+)] II; lon-1(e185) III* |
| **GCP1175** | *prtSi222[Pwrt-2::GFP::his-58::GFP::PH(PLC1δ1); cb-unc-119(+)] II; unc-119(ed3) III; dpy-11(e224) V* |
| **GCP1177** | *ItSi249[PdIg-1delta7::dlg-1::GFP; cb-unc-119(+)] I; prtSi239[Pwrt-2::mCherry::his-58::mCherry::PH(PLC1δ1); cb-unc-119(+)] II; unc-119(ed3) III; dyci-1(prt152[Δ4-36]) IV/nT1[qIs51] (IV;V)* |
| **GCP1188** | *prtSi222[Pwrt-2::GFP::his-58::GFP::PH(PLC1δ1); cb-unc-119(+)] II; lon-1(e185) III; dyci-1(prt152[Δ4-36]) IV/nT1[qIs51] (IV;V)* |
| **GCP1205** | *dhc-1(he250[mCherry::dhc-1]) I; lin-5(cp288[lin-5::mNG-C1<sup>3xflag</sup>]) II* |
| **GCP1287** | *prtSi222[Pwrt-2::GFP::his-58::GFP::PH(PLC1δ1); cb-unc-119(+)] II; unc-119(ed3) III; dyci-1(prt152[Δ4-36]) IV/nT1[qIs51] (IV;V); dpy-11(e224) V/nT1[qIs51] (IV;V)* |
| **GCP1325** | *prtSi222[Pwrt-2::GFP::his-58::GFP::PH(PLC1δ1); cb-unc-119(+)] II; ani-1(syb4317 [TagRFP::ani-1]) III* |
| **GCP1341** | *prtSi222[Pwrt-2::GFP::his-58::GFP::PH(PLC1δ1); cb-unc-119(+)] II; ani-1(syb4317 [TagRFP::ani-1]) III; dyci-1(prt152[Δ4-36]) IV/nT1[qIs51] (IV;V)* |
| **GCP1430** | *ItSi570[Pdpy-7::GFP::tbb-2::mCherry::his-11; cb-unc-119(+)] I; lin-5(ev571) II; unc-119(ed3) III* |
| **GCP1431** | *goa-1(vsSi32[goa-1::GFP]) I; prtSi239[Pwrt-2::mCherry::his-58::mCherry::PH(PLC1δ1); cb-unc-119(+)] II; unc-119(ed3) III* |
| **GCP1438** | *apr-1(cp166[mNG-C1^3xFLAG::apr-1]) I; prtSi239[Pwrt-2::mCherry::his-58::mCherry::PH(PLC1δ1); cb-unc-119(+)] II; unc-119(ed3) III* |
| **GCP1463** | *prtSi239[Pwrt-2::mCherry::his-58::mCherry::PH(PLC1δ1); cb-unc-119(+)] II; unc-119(ed3) III; prtSi262[Pscm::NLS(egl-13)::GFP::NLS(SV40)::GFP:PH(PLC1δ1); cb-unc119(+)] V* |
| **GCP1464** | *prtSi239[Pwrt-2::mCherry::his-58::mCherry::PH(PLC1δ1); cb-unc-119(+)] II; unc-119(ed3) III; dyci-1(prt152[Δ4-36]) IV/nT1[qIs51] (IV;V); prtSi262[Pscm::NLS(egl-13)::GFP::NLS(SV40)::GFP:PH(PLC1δ1); cb-unc119(+)] V/nT1[qIs51] (IV;V)* |
| **GCP1468** | *apr-1(cp166[mNG-C1^3xFLAG::apr-1]) I; prtSi239[Pwrt-2::mCherry::his-58::mCherry::PH(PLC1δ1); cb-unc-119(+)] II; unc-119(ed3) III; dyci-1(prt152[Δ4-36]) IV/nT1[qIs51] (IV;V)* |

